# Detailed *in vivo* characterization of the players and impact of individual loop extrusion trajectories

**DOI:** 10.1101/2023.01.04.522689

**Authors:** Ruiqi Han, Yike Huang, Michelle Robers, Mikhail Magnitov, Iwan Vaandrager, Amin Allahyar, Marjon J.A.M. Verstegen, Kexin Zhang, Elzo de Wit, Wouter de Laat, Peter H.L. Krijger

**Affiliations:** Oncode Institute, Hubrecht Institute-KNAW and University Medical Center Utrecht, Utrecht, the Netherlands; Division of Gene Regulation, The Netherlands Cancer Institute, Amsterdam, the Netherlands

## Abstract

The cohesin complex shapes chromosomes by DNA loop extrusion, but individual extrusion trajectories were so far unappreciable *in vivo*. Here, we developed and validated TArgeted Cohesin Loader (TACL), a system enabling the strong activation of anchored loop extrusion from dozens of defined genomic sites in living cells. Studying their individual loop extrusion trajectories revealed that extruding cohesin^STAG2^ stops not only at domain boundaries but at all flanking CTCF sites, engaging them in a complex transient looping network that supports intradomain contacts. Cohesin^STAG1^ cannot associate with weak CTCF binding sites and fails to similarly support intradomain interactions. NIPBL-MAU2 remains associated with cohesin when stalled at looping CTCF sites, suggesting these factors may also be required for loop stabilization. TACL induces cohesin traffic jams and illegal loops with divergent CTCF sites, demonstrating that stalled cohesin can block extruding cohesin *in vivo*. Genes exposed to TACL-induced loop extrusion were collectively hindered in transcription and the underlying chromatin altered its accessibility and reduced its H3K27ac marks. Thus, by enabling the study of individual loop extrusion trajectories *in vivo,* we could assign new functions to players, identify new looping networks and uncover an interplay between loop extrusion, gene transcription and chromatin composition.

## Introduction

The evolutionary conserved cohesin complex is a tripartite ring-shaped structure consisting of RAD21, SMC1 and SMC3, associated with either STAG1 or STAG2. The complex functions to hold sister chromatids together during mitosis and to shape chromosomes in interphase cells by forming topologically associating domains (TADs)^1–3^. Cohesin establishes TADs presumably by loop extrusion^4–6^ in which the complex is loaded on chromatin and subsequently reels in flanking sequences to build progressively larger DNA loops in an ATP-dependent manner^7–10^. STAG1 and -2 have overlapping but non-redundant functions. STAG1-associated cohesin is more stably associated with chromatin and crucial to establish the longer chromatin loops between CTCF-associated domain boundaries. STAG2-associated cohesin was reported to be more often found at non-CTCF sites and form intra-TAD loops between enhancers and genes^11–13^. Cohesin cycles between a chromatin-bound and -extruding state and an unbound state^14,15^. It requires NIPBL and MAU2 for stable chromatin association^16,17^ and loop extrusion processivity^18–21^, while WAPL releases cohesin from chromatin and restricts loop extrusion^22^. When reaching convergently oriented CTCF proteins that demarcate domain boundaries, cohesin is protected against WAPL-mediated release^22^: it stalls and forms temporarily stabilized chromatin loops between opposite domain boundaries^15,23–25^. PDS5 interacts with cohesin and localizes on chromosomes at CTCF-bound loop anchors^3,26,27^. It is thought to compete with NIPBL for binding to cohesin^28,29^ and, similar to CTCF, PDS5 functions to restrict chromatin loop sizes^3,26,30^. How loop extrusion impacts transcription remains unclear, recent evidence suggests that the transcription and loop extrusion machineries interact when traversing chromatin^31–33^ and that continuous cohesin-mediated loop extrusion is required for the regulation of developmental genes by distal enhancers^14,34–37^. Studying the impact of cohesin loop extrusion activity *in vivo* remains challenging though^38^, as individual loop extrusion trajectories cannot be manipulated in living cells. Furthermore, *in vivo* studies of cohesin largely rely on (acute) cohesin protein depletion, which leads to widespread changes in chromatin structure and functioning, cell cycle arrest and cell death^1,14,15^. Systems that enable local control of cohesin loop extrusion activity at defined genomic locations in healthy living cells will help to more accurately monitor the direct consequences of altered loop extrusion activity.

## Results

### TArgeted Cohesin Loader (TACL) system

Here we developed the TArgeted Cohesin Loader (TACL) system, a genetic platform for site-specific initiation and manipulation of individual loop extrusion trajectories *in vivo*. TACL employs the TetO/TetR system, introducing the TetR peptide fused to the cohesin loading factor MAU2 (SCC4) to conditionally recruit cohesin and initiate loop extrusion trajectories from Tet operator sequences integrated in the human genome. We utilized the PiggyBac transposon system^39^ to create a human eHAP1 cell line with twenty-seven TetO platforms (hereafter “TetO”) randomly inserted across 19 different chromosomes (Fig.1A, B). Each TetO platform contained 48 TetR binding sites. We chose eHAP1 cells for our studies as they are haploid and therefore have no untargeted chromosome copies compromising the monitoring of TACL-induced chromosomal changes. Cells were transduced with lentivirus to stably express TetR fused to FLAG-MAU2 (“TACL”) or to FLAG-mCherry (“Cherry”). Western blotting showed that TetR-FLAG-MAU2 mainly localized to the cytoplasm, but also entered the nucleus, where it replaced most (∼85%) endogenous MAU2 protein (fig. S1A). This was not unexpected as others previously found that MAU2 and NIPBL interact with each other to maintain their mutual stability in the nucleus^22,40,41^. TetR-MAU2 therefore competed with endogenous MAU2 for binding to NIPBL. We assessed whether TetR-MAU2 replacing endogenous MAU2 in the nucleus had any general impact on cohesin distribution, chromosome topology or gene expression in TACL cells. To this end, we studied genomic intervals that were at least 3Mb away from TetO integration sites. ChIP-seq for SMC1 and RAD21 showed that the distribution of cohesin along chromosomes was similar between TACL cells and control Cherry cells (Fig. S1B). Hi-C demonstrated that also chromatin topology, i.e. chromatin loops, TAD boundaries and TADs, was not altered in a meaningful way (Fig. S1C, D). With nascent RNA-seq we observed only a few genes that were differentially expressed between TACL and Cherry cells (Fig. S1E). Finally, the cellular proliferation rates of TACL and Cherry cells were comparable (Fig. S1F). Therefore, we concluded that TetR-MAU2 was capable of functionally replacing endogenous MAU2 in eHAP1 cells.

**Figure 1.**
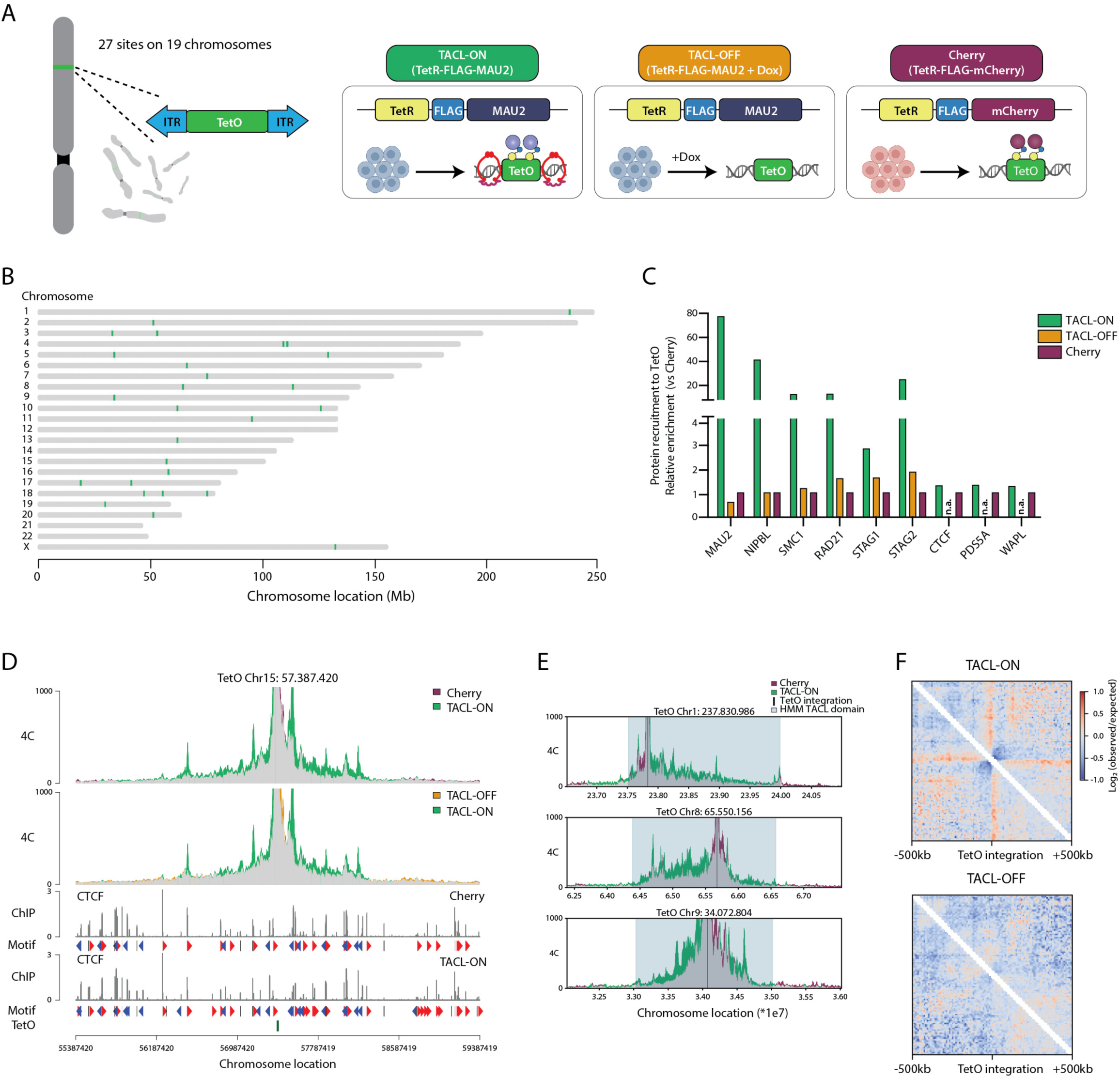
TACL recruits functional cohesin complex at designated genomic sites. **(A)** Cartoon illustration of the TACL system. Three experimental conditions are indicated in different colors as ‘TACL-ON’, ‘TACL-OFF’, and ‘Cherry’. TACL-ON and TACL-OFF cells (blue) were transduced with lentivirus expressing TetR-FLAG-MAU2 and Cherry cells (pink) were transduced with lentivirus expressing TetR-FLAG-mCherry. **(B)** Chromosomal distribution of the TetO platforms in eHap1 cells. Inserted TetO platforms are illustrated in green. **(C)** Relative ChIP-seq enrichment of the indicated proteins at the TetO platforms. Values are normalized to Cherry condition. n.a. indicates data not available for this condition. **(D)** 4C overlay and CTCF ChIP-seq tracks of an example locus of the TetO platform. The plot is centered at the viewpoint of the 4C analysis, which is also the location of the TetO platform (indicated at the lower part of the panel). Common (no difference) interactions are indicated in gray. **(E)** Examples of the Hidden Markov Model (HMM) TACL domains annotated based on the 4C-seq profiles of TACL-ON and Cherry cells. TetO platform is marked in black and HMM domain is highlighted in light blue. Common (no difference) interactions are indicated in gray. **(F)** Average interactions centered at TetO integrations. Note the stripes emerging from TetO platforms.

### TACL recruits the cohesin complex to defined genomic locations

Next, we investigated the consequences of TetR-MAU2 recruitment to the genomically integrated TetO sites. ChIP-qPCR with primers amplifying TetO sequences allowed the analysis of TetR-MAU2 recruitment to all TetO sites at once. Because all TetO sites are identical in sequence, studying recruitment to individually integrated TetO sites was not possible. We confirmed that the TetR-MAU2 and TetR-mCherry fusion proteins efficiently bound to TetO (Fig. S2A). TetR binding to TetO was doxycycline (Dox) reversible: the TetR fusion proteins were released from TetO when cells were treated with Dox for one hour (Fig. S2A). TetR-MAU2, not TetR-mCherry, selectively co-recruited NIPBL to TetO and attracted the core cohesin subunits SMC1 and RAD21, as seen before in yeast^42^. Similarly, when we added Dox for one hour, these factors were no longer recruited (Fig. S2A). We referred to the bound condition (no Dox) as “TACL-ON”, and to the unbound condition (+ Dox, 1 hour) as “TACL-OFF”. The TetR-mCherry (Cherry) cells served as an alternative negative control condition.

We then used ChIP-seq experiments to further analyze which factors were co-recruited by MAU2 to TetO. When we pulled down chromatin associated with FLAG (-tagged TetR-MAU2), MAU2, NIPBL, SMC1, and RAD21 (Fig. 1C), strong enrichments of TetO sequences (over genomic sequences) were observed in TACL-ON cells, but not in TACL-OFF or Cherry cells, as also seen with ChIP-qPCR (Fig. S2A). ChIP-seq with antibodies against the cohesin subunits STAG1 and STAG2 showed that mainly STAG2 was co-loaded onto TetO (Fig. 1C). This further confirmed that the targeting of MAU2 to TetO triggered the co-recruitment of NIPBL and core cohesin subunits. CTCF, PDS5A and WAPL, as well as cohesin-associating proteins, were not recruited by MAU2 to TetO (Fig. 1C).

### TACL enables controlled local activation of cohesin loop extrusion

To test whether TACL enabled controlled activation of cohesin loop extrusion from TetO sites, we first performed 4C-seq to investigate changes in chromatin contacts made by the integrated TetO cassettes. Since one can assume that each TetO predominantly contacted its own linear surrounding sequences, per condition a single pair of 4C primers could be used to simultaneously assess the individual contact profiles of all 27 TetO locations. We observed that all TetO sites engaged in more long-range contacts (>200 kb) at the expense of short-range contacts in the TACL-ON condition (fig. S2B). Clearly noticeable was that TACL induced many of the TetO platforms to form strong specific interactions with surrounding CTCF sites, often over hundreds of kilobases (Fig. 1D, and fig. S2C). The increased long-range contacts were absent in Cherry cells and were dismantled in TACL-OFF cells (Fig. 1D, E, S2C). This strongly suggested that TACL enables controlled activation of chromatin loop formation from integrated TetO sites in living cells.

The gain in 4C signal in TACL-ON versus Cherry cells was used in a Hidden Markov Model (HMM) to define the domains with TACL-induced looping trajectories (TACL-domains) (Fig. 1E, see methods). These domains often extended asymmetrically from a TetO site and spanned an average genomic distance of 2.56 Mb (minimum 1.08 Mb, maximum 4.43 Mb) (Fig. 1E and Fig. S2D). The distances to boundaries, we assumed, reflected the maximum lengths of TACL-induced cohesin loop extrusion trajectories.

### Architectural stripes suggest that TACL induces TetO-anchored loop extrusion events

TACL-induced local 3D genome changes were further confirmed by Hi-C analysis. By averaging the chromatin contact data around all TetO sites, stripes, not chromatin jets, were seen emerging in a bi-directional manner from the TetO sites (Fig. 1F and Fig. S2E). Chromatin jets describe a recently identified Hi-C signature observed at some locally dominant cohesin loading sites. They reflect bouquets of unanchored, bi-directionally extruding cohesin molecules that all initiated extrusion at the same site^29^. In contrast, stripes, normally only observed at strong CTCF boundaries^15,43^, are believed to reflect differently sized chromatin loops formed by uni-directional extruding cohesin molecules anchored at these sites. The observation that TetO platforms also functioned as sites holding cohesin for single-sided loop extrusion activity may be explained by the high number of cohesin molecules constantly present on the 48xTetO platforms or the movement of cohesin is constrained by tethering to TetR-FLAG-MAU2. If they interfere with each other’s extrusion activity, only the outer cohesin molecules may be able to reel in their respective flanking sequences, seen as bi-directional stripes in Hi-C.

### Defining the ends of TACL-induced loop extrusion trajectories

To understand what happens to cohesin and its interaction partners in TACL domains we performed a series of ChIP-seq experiments. Across the genome of TACL cells FLAG-tagged MAU2 is typically associated with active enhancers and promoters, as also seen for endogenous MAU2 in Cherry cells (see below), in a manner that was insensitive to Dox treatment. However, inside the TACL domains FLAG-tagged MAU2 accumulated at many novel sites and here MAU2 deposition was dependent on the active TACL system (Fig. 2A). When we selected these sites, we observed that they also accumulated NIPBL, SMC1, RAD21, STAG1 and STAG2, in a TACL-dependent manner (Fig. 2B-D). In addition, we found that TACL-induced factor deposition occurred at CTCF sites that also recruited PDS5A and WAPL (Fig. 2C).

**Figure 2.**
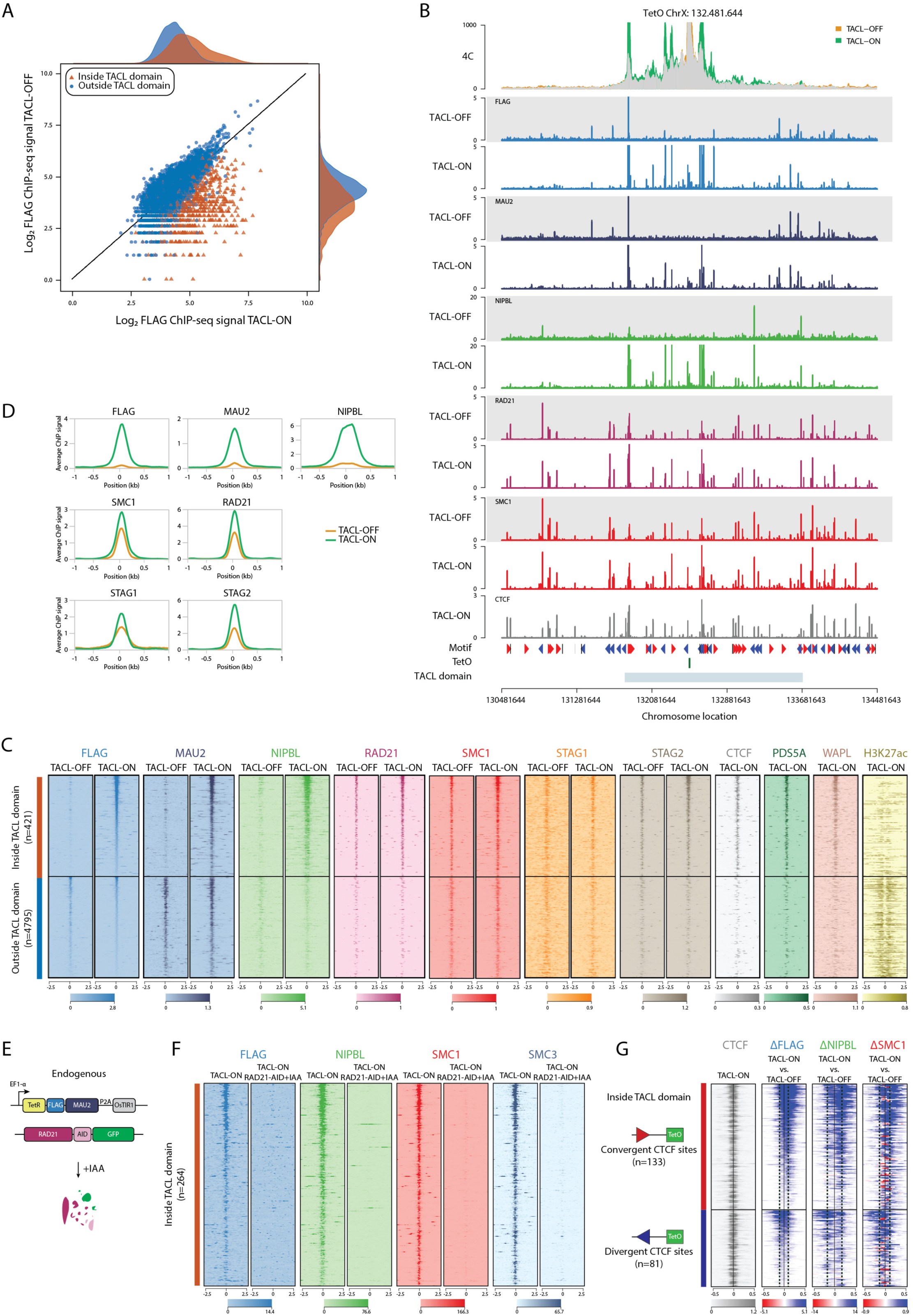
Local accumulation of cohesin complex reveals important features of individual subunits. **(A)** Log2 fold change of FLAG ChIP-seq signal in TACL-ON and TACL-OFF conditions. FLAG peaks inside or outside the TACL domain are depicted in red or blue respectively. The Dox-responsive FLAG peaks are mostly located within the TACL domain. **(B)** 4C overlay and ChIP-seq tracks of an example locus of the TetO platform. TACL-OFF tracks (grey) serve as controls. The plot is centered at the viewpoint of the 4C analysis (indicated as the TetO platform at the lower part of the panel). TACL domain indicates the loop extrusion domain defined by HMM model. **(C)** Heatmaps of ChIP-seq signals for the cohesin-related factors. FLAG peaks are plotted at the center ± a 2.5kb region. The color map at the bottom shows an increased in signal from light to dark color intensities. Upper half (red bar on the left) shows the signals at differential FLAG peaks within the TACL domain; lower half (blue bar on the left) shows the signals at FLAG peaks outside the TACL domain. ‘n’ stands for the number of peaks. **(D)** Histogram profiles of ChIP-seq signals for different cohesin subunits at local differential FLAG peaks. Signals are scaled to the averag coverage outside the TACL-domains. **(E)** An illustration of the RAD21 degron system. **(F)** Heatmaps of ChIP-seq signals for FLAG, NIPBL, SMC1, and SMC3 in RAD21-AID TACL-ON cells. Peaks shown are differential FLAG peaks at CTCF sites inside the TACL domain. **(G)** Heatmaps of differential ChIP-seq signals for FLAG, NIPBL, and SMC1 at surrounding CTCF sites between TACL-ON and TACL-OFF conditions. Peaks shown are CTCF peaks inside the TACL domain overlapping with a differential FLAG peak. Upper half (red bar on the left) shows the CTCF sites with a convergent orientation towards the TetO platform; lower half (blue bar on the left) shows the CTCF sites with a divergent orientation against the TetO platform. ‘n’ stands for the number of CTCF sites. The color map at the bottom shows an increased signal in TACL-ON cells in blue and decreased signal in red.

To investigate if the deposition of MAU2 and NIPBL at flanking CTCF sites was the consequence of their concentration at TetO sites and subsequent nucleoplasmic diffusion, or of their co-migration with cohesin extruding from TetO, we knocked-in an auxin-inducible degron (AID2) ^44^ to acutely deplete endogenous RAD21 in the TACL and Cherry cells (Fig. 2E). Treating the TACL cells for two hours with 5-Ph-IAA (IAA) degraded RAD21 to undetectable levels (fig. S3A) and dismantled the TACL-induced topological contacts, as determined by 4C-seq (Fig. S3B). TetR-MAU2 and NIPBL were still efficiently recruited to TetO (Fig. S3C), but could not be detected at the flanking CTCF sites, where SMC1 and SMC3 were also no longer found (Fig. 2F). We therefore conclude that MAU2 and NIPBL accumulated at flanking CTCF sites through co-migration with and subsequent pausing of TACL-induced loop extruding cohesin complexes. The colocalization of WAPL and PDS5A at these CTCF sites^29^ suggested that they often served as the chromatin-unloading sites of cohesin, thus marking the ends of loop extrusion trajectories initiated from TetO.

### Cohesin traffic jams at blocking CTCF sites

Notably, not all CTCF sites in the TACL domains were detectably used as pause or end points of TACL-induced loop extrusion trajectories. This could partially be explained by differences in CTCF’s occupancy at these sites, and their chromosomal distance from TetO (Fig. S4A). Given that CTCF-binding orientation on DNA is important for halting cohesin loop extrusion^23–25^, we examined whether the blocking sites were facing towards (convergent) or away (divergent) from TetO. As expected, it was predominantly the CTCF sites facing TetO that most effectively blocked TACL-induced loop extrusion (Fig. S4A). Interestingly though, divergent CTCF sites (looking away from TetO) also accumulated cohesin as a consequence of TACL (Fig.2G). Detailed inspection revealed that TACL-induced cohesin deposition at convergently oriented CTCF sites led to queuing of multiple extruding cohesin holoenzymes in front of, but also behind the CTCF sites. Similar queues were also present in front of and behind divergently oriented (‘illegal’) CTCF molecules (Fig. 2G). *In vitro*, purified condensin complexes were reported to pass each other when extruding naked DNA^45^. Our data however suggested that *in vivo*, at chromosomal locations with high loop extrusion activity, extruding cohesin complexes can be stalled when encountering another cohesin complex bound to a CTCF roadblock. Cohesin ‘traffic jams’ were previously suggested to exist *in vivo* based on the observation of loop collision events^46–48^; our ChIP-seq results provided direct evidence for this.

### V5-MAU2 molecules do not replace TetR-MAU2 on cohesin extruding from TetO

Given that NIPBL and MAU2 were not known to associate with CTCF sites, we were surprised to find that loop extrusion initiation from TetO resulted in high accumulation of NIPBL and MAU2 at the flanking CTCF sites. When we called peaks in our NIPBL and MAU2 ChIP-seq datasets from control Cherry cells, they were typically located at active promoters and enhancers, not at CTCF sites (Fig. S4B). However, when we selected and combined all SMC1 and NIPBL binding sites and stratified them according to co-localization with CTCF sites, enhancers or promoters, we found NIPBL and MAU2 also enriched at CTCF sites across the genomes of both TACL and (Fig. 3A) control Cherry cells (Fig. S4C). Moreover, when we depleted RAD21 from TACL and control Cherry cells, NIPBL and MAU2 remained associated with enhancers and promoters, but disappeared from CTCF sites (Fig. 3A and Fig. S4C). This demonstrated that also in wildtype cells NIPBL and MAU2 are brought by cohesin to CTCF sites. Close inspection of a published dataset confirmed that others found NIPBL associated with CTCF sites^12^.

**Figure 3.**
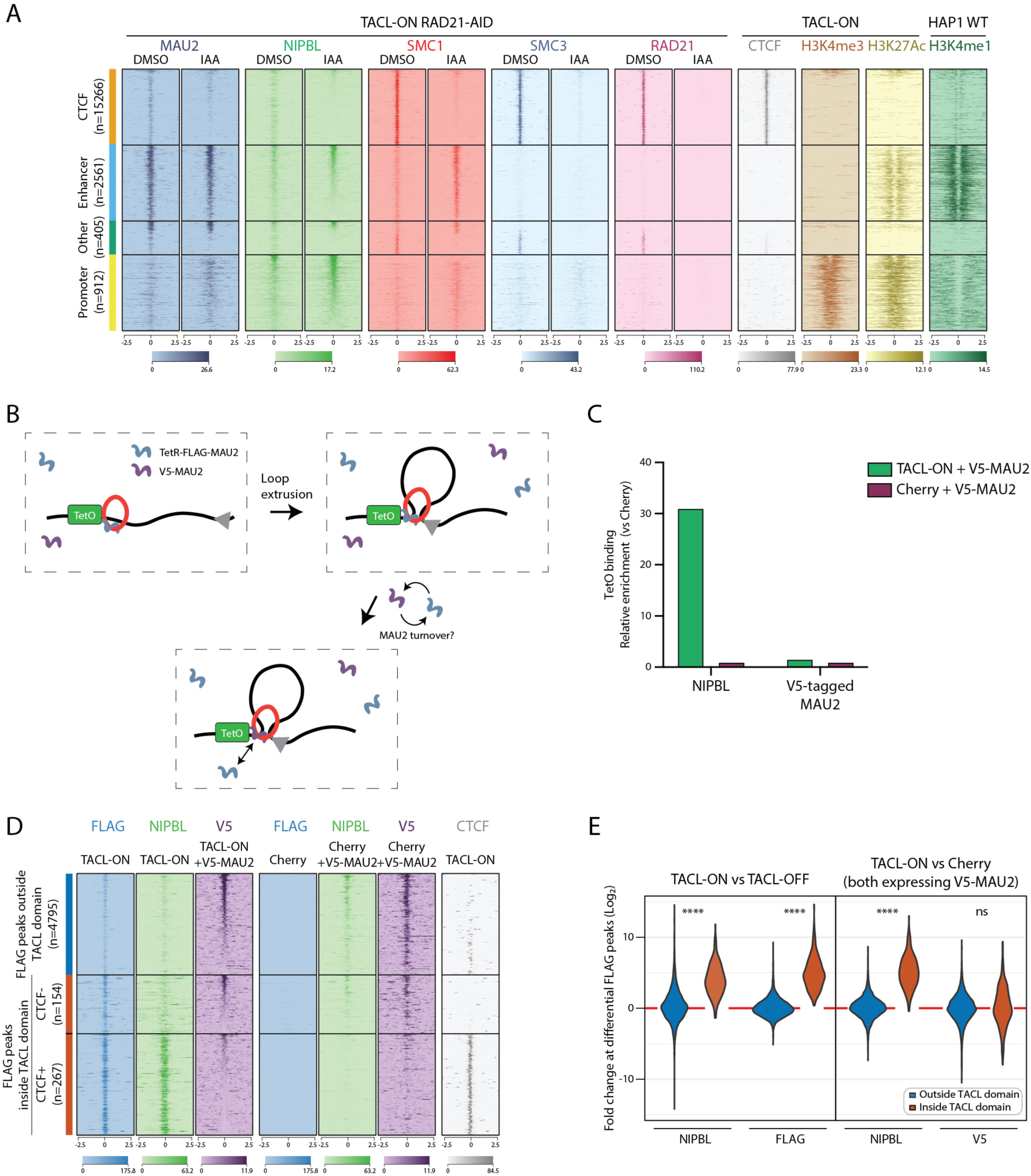
MAU2 forms a stable extruding complex with cohesin. **(A)** Heatmaps of ChIP-seq signals for MAU2, NIPBL, SMC1, SMC3, and RAD21 in TACL-ON RAD21-AID cells. For reference, CTCF, H3K4me3, H3K27Ac, and H3K4me1 signals are displayed. SMC1 and NIPBL peaks located outside TACL-domains and > 3MB from the TetO integration side are categorized into four groups as indicated on the left, CTCF, Enhancer (H3K4me3-/H3K27Ac+ and/or H3K4me1+), Other (no active histone modification marks), and Promoter (H3K4me3+). ‘n’ stands for the number of peaks. **(B)** A cartoon illustration of the TetR-FLAG-MAU2 and V5-MAU2 co-expressing experiment. ChIP-seq signals of either FLAG or V5 represents the enrichment of the different tagged-MAU2 on chromatin. **(C)** Relative enrichment of NIPBL and V5-MAU2 at the TetO platforms. Values are normalized to Cherry condition. **(D)** Heatmaps of ChIP-seq signals for FLAG, NIPBL, and V5 in TACL-ON and Cherry cells co-expressing V5-MAU2. FLAG peaks in TACL-ON cells are divided into three categories (shown on the left), outside TACL domain, inside TACL domain with CTCF binding, and inside TACL domain without CTCF binding. ‘n’ stands for the number of FLAG peaks. **(E)** Violin plots of log2 fold change at FLAG peaks outside TACL domains and differential FLAG peaks within TACL domains located at CTCF binding sites in TACL-ON cells. The plots are divided into two groups per factor based on the location of the FLAG peaks relative to the TACL domain. Left panel represents the comparison of NIPBL and FLAG signals between TACL-ON and TACL-OFF and the right panel represents the comparison of NIPBL and V5 signals between TACL-ON and Cherry cells co-expressing V5-MAU2. Note the similar levels of V5 signals inside and outside the TACL domain compared to other factors. Statistical significance determined using the Mann-Whitney U-test: **** p < 0.0001; ‘ns’ indicates not significant.

Having confirmed that extruding cohesin can transport NIPBL-MAU2 to CTCF sites, we investigated turn-over of MAU2 association with extruding cohesin. Previous *in vitro* experiments suggested that the extruding cohesin complex transiently associated with NIPBL, and presumably MAU2, and that a dynamic exchange of NIPBL (and MAU2) was needed for cohesin to act as an active loop-extruding holo-enzyme^19,20^. To analyze *in vivo* whether NIPBL and MAU2 stably or transiently associated with cohesin complexes traveling along chromosomes, we co-expressed V5-tagged MAU2 with FLAG-tagged TetR-MAU2 in TACL cells (Fig. 3B). Together, the two MAU2 fusion proteins replaced all endogenous MAU2 protein (Fig S4D). Elsewhere in the genome (outside of TACL domains and >3Mb away from TetO), V5-MAU2 occupied the same sites as MAU2 (and TetR-MAU2), being mostly active promoters and enhancers but also weakly at CTCF sites that required RAD21 for MAU2 recruitment (fig S4E). This confirmed that V5-MAU2 behaved similarly to endogenous MAU2: it associated with the chromatin entry sites of cohesin and was transported by cohesin to CTCF sites. V5-MAU2 has no affinity for and is not recruited to TetO though (Fig 3C) and it also remains absent from the TetO-flanking CTCF sites (Fig. 3D, E). Thus, while constant NIPBL exchange appears necessary for cohesin to continue loop extrusion *in vitro* ^19,20^, we found no evidence for MAU2 molecules replacing each other on cohesin extruding from TetO *in vivo*.

### Factors controlling the lengths of loop extrusion trajectories

Hi-C experiments showed that the depletion of WAPL^22^, PDS5^49^ and STAG2^11^ resulted in CTCF-CTCF contacts over larger distances. This is usually interpreted as evidence for extended loop extrusion trajectories, but formal evidence for this requires studying individual extrusion trajectories and their length. With 25 TetO integration sites located in separate TACL domains, the TACL system enabled direct monitoring of 50 cohesin loop extrusion trajectories from start to end. We created additional inducible degron lines for WAPL, PDS5A, STAG2 and CTCF (Fig. 4A, B). Depletion of CTCF has been implicated to lead to unimpeded extrusion by cohesin^50^. The 4C profiles confirmed this: in the absence of CTCF, TetO sites no longer formed specific loops with CTCF sites but instead engaged in more distal, more uniformly distributed chromatin contacts (Fig. 4C, D ‘CTCF depleted’). Although WAPL, PDS5A and STAG2 depletion does not impact CTCF binding to chromatin^22^, their depletion was reported to result in extended chromatin loops. Our data confirmed this at the level of individual loop extrusion trajectories. In the absence of the cohesin-unloading factor WAPL^22^ in TACL-ON cells, the TetO sites no longer engaged in frequent contacts with proximal CTCF sites, but instead formed prominent interactions with more distal CTCF sites, sometimes over 4-5 megabases (Fig. 4C, D, ‘WAPL depletion’). When we depleted PDS5A, which restricts chromatin loop sizes by competing with and displacing NIPBL from cohesin^3,26,30^, or STAG2^11–13^, which causes cohesin to less stably associate with chromatin than STAG1 cohesin^STAG1^, we found similar chromatin contact changes: original TACL-induced preferred contacts between TetO and flanking CTCF sites were weakened and new specific contacts with more distal CTCF sites were established. The new contact partners of TetO were very similar between the WAPL- and PDS5A- and STAG2-depleted cell lines and also the TACL-induced loop extrusion trajectory lengths increased similarly, creating much larger TACL domains as determined by HMM from the differential 4C contacts maps. While originally the TACL domains measured an average genomic distance of 2.55 Mb, the average domain size increased to 6.39 Mb (max 8.23 Mb), 5.91 Mb (max 7.98 Mb) and 5.59 Mb (max 8,39 Mb) respectively in WAPL-, STAG2- and PDS5A-depleted cells (fig. S5A, B).

**Figure 4.**
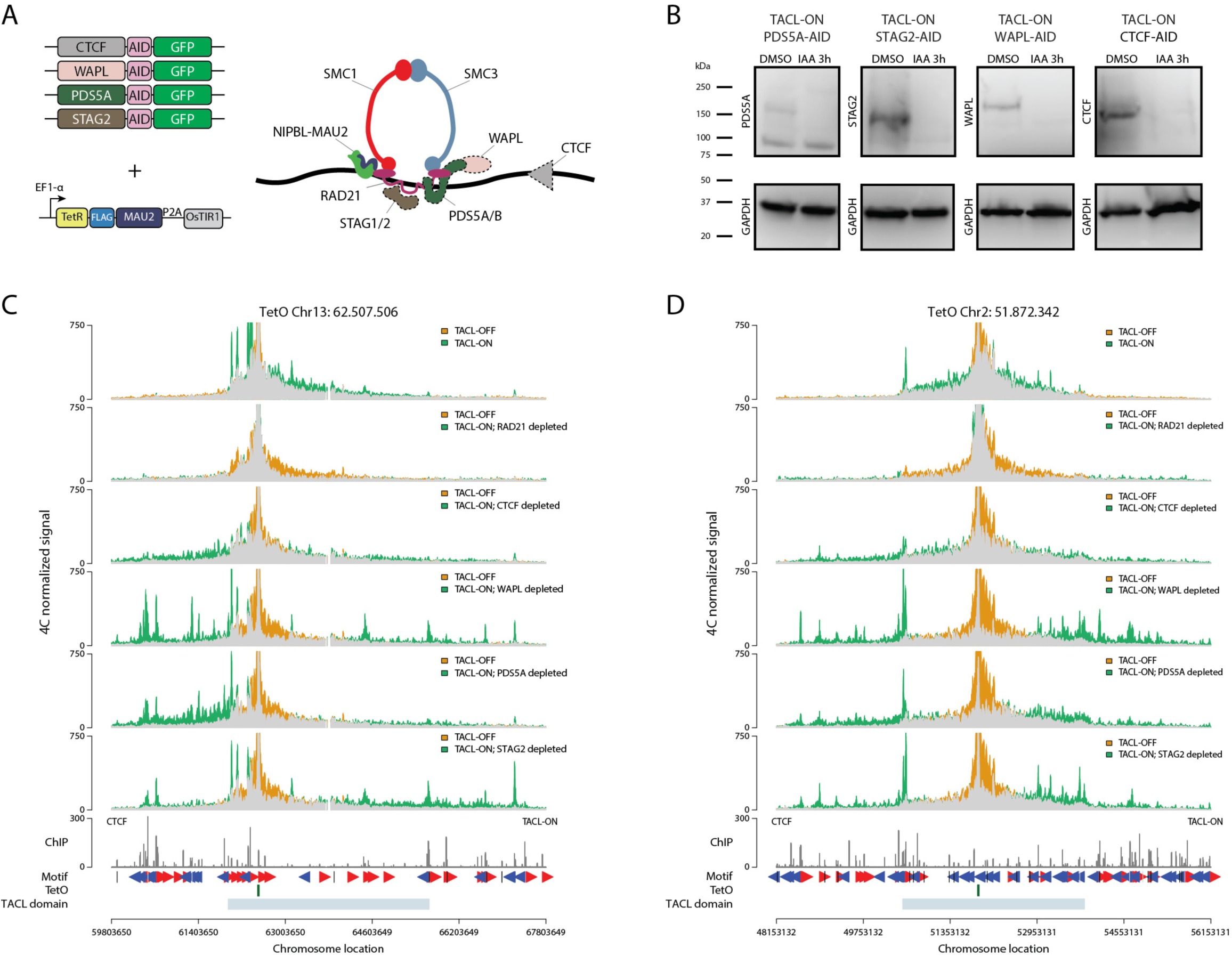
Depletion of cohesin factors causes loop extension. **(A)** Cartoon illustration of the inducible degron systems of PDS5A, STAG2, WAPL, and CTCF. **(B)** Western blot images of the degrons of the cohesin factors. Comparisons were made between DMSO or 3hours of IAA treatment. Proteins were detected using the corresponding antibodies. GAPDH was used as loading control. **(C-D)** 4C tracks of two example TetO sites. Comparison was made between either TACL-OFF and TACL-ON or TACL-ON after protein depletion in the corresponding degron cell lines.

### Cohesin^STAG1^ can replace cohesin^STAG2^ at some but not all intradomain CTCF sites

Next, we focused on the STAG proteins and used TACL to gain further insights into the roles of STAG1 and -2 in the topological organization of chromatin. With 4C-seq we observed that without STAG2, cohesin^STAG1^ engaged TetO in new specific chromatin loops with even more distal CTCF sites, sometimes almost 5 Mb away (Fig. 5A). Simultaneously, we noticed a dramatic collapse of local intradomain contacts at all TACL domains in STAG2-depleted cells (Fig. 5A and S5A, note the loss of local contacts in the grey common region in STAG2-depleted cells). This suggested that only cohesin^STAG2^, not cohesin^STAG1^, was capable of supporting these local intradomain contacts. To investigate this further, we first used the 4C-seq contact profiles and HMM to systematically define the domains with collapsed interactions in STAG2-depleted cells (“collapsed domains”) (fig. S5C). We then generated ChIP-seq data for FLAG, NIPBL, STAG1 and SMC1 in STAG2-depleted TACL-ON cells, and compared their binding sites to those previously observed in TACL-ON cells. We identified three categories of CTCF sites surrounding TetO (n = 609) that all recruited cohesin^STAG2^ but having a distinct capacity to also halt cohesin^STAG1^. Category I sites (n=164), the ‘shared STAG1/2’ sites, recruited both cohesin^STAG2^ and cohesin^STAG1^ in wildtype cells. Category II sites (n=245), the ‘conditional STAG1’ sites, recruited cohesin^STAG1^ only in STAG2-depleted cells. Category III sites (n=200), the ‘STAG2-only’ sites, never recruited cohesin^STAG1^, also not in STAG2-depleted cells (Fig. 5B). The three types of CTCF sites seemed randomly distributed along the TetO-flanking regions (Fig. 5A). Category I sites were typically the strongest CTCF sites (Fig. 5C) that also accumulated most STAG2 (fig. S5D) and cohesin (fig. S5E). Oppositely, category III sites marked the weakest CTCF binding sites, with least STAG2 and cohesin recruitment (Fig. 5C, D). Thus, cohesin^STAG2^ appeared superior to cohesin^STAG1^ in associating with the weakly DNA-bound CTCF molecules.

**Figure 5.**
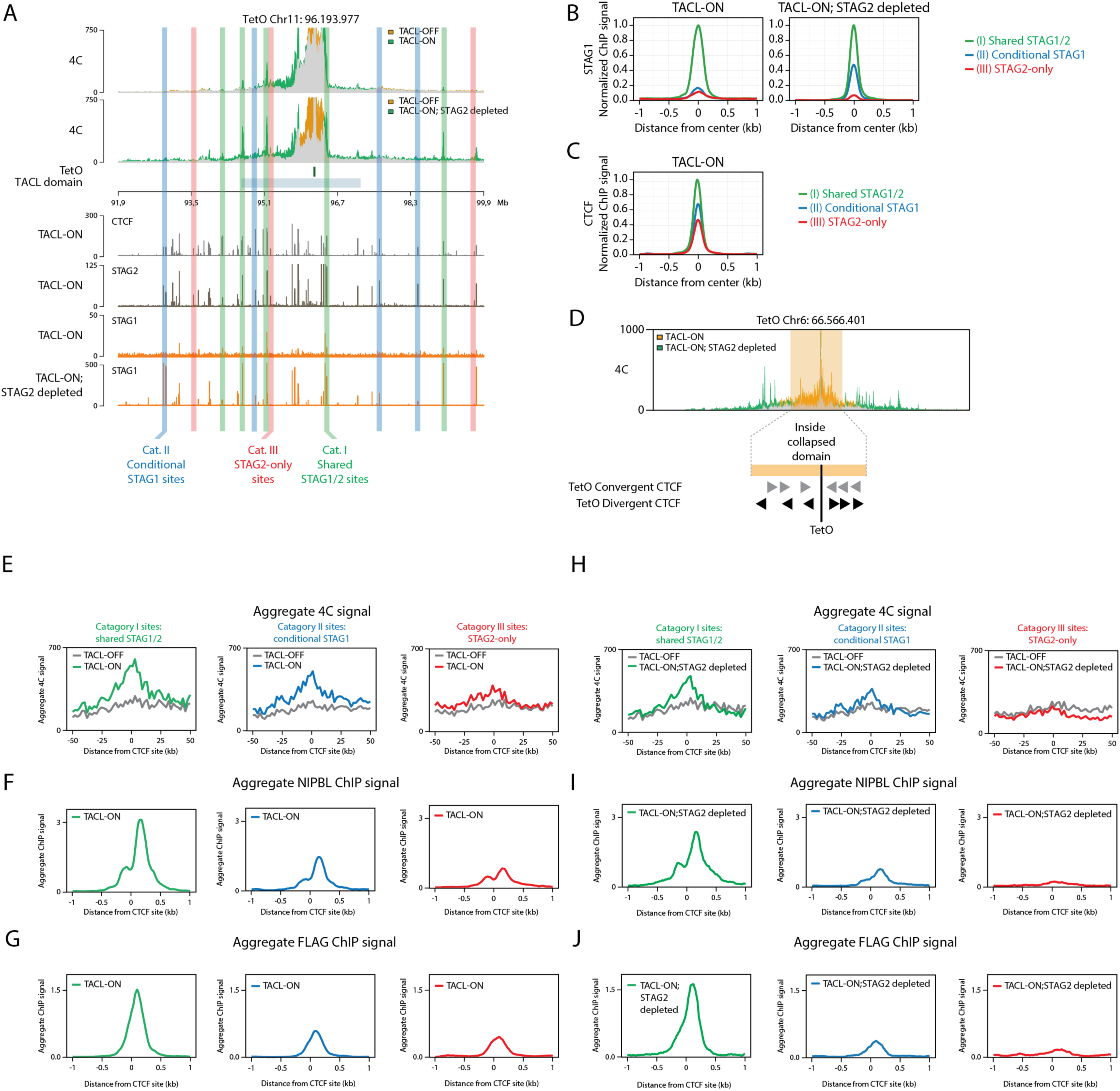
STAG2 depletion reveals different roles of STAG1 and STAG2 on chromatin. **(A)** 4C and ChIP-seq tracks of an example TetO site in TACL-ON and TACL-ON, STAG2-depleted cells. Different categories of STAG sites are highlighted. Category I shared STAG1/2 site is marked in green; category II conditional STAG1 site is marked in blue; category III STAG2-only site is marked in red. Only a selection of examples of each category are indicated in the panel. **(B)** Average signal plots of STAG1 ChIP-seq signals at STAG1/2**-**CTCF sites within the TACL domain. Values are normalized to shared SA1/2 for comparisons. Each category is color-coded as illustrated in the panel. **(C)** Average signal plots of CTCF ChIP-seq signals at STAG1/2 and CTCF sites within the TACL domain. Values are normalized to shared SA1/2 for comparisons. Each category is color-coded as illustrated in the panel. **(D)** An illustration of the collapsed intradomain in STAG2-depleted cells (highlighted in orange). 4C overlay of TACL-ON and TACL-ON, STAG2-depleted conditions are shown. In the lower panel, an illustration of TetO-convergent and TetO-divergent sites is shown. **(E)** Aggregate 4C signal plots of TACL-ON and TACL-OFF cells within the STAG2-depleted collapsed domain. Only CTCF sites with TetO-convergent orientations are shown. Panels are divided into different STAG categories. TACL-OFF is indicated in grey and TACL-ON is indicated with the colors of each STAG category. Cat. I in green, cat. II in blue and cat. III in red. Signals are centered around CTCF binding sites. **(F)** Aggregate NIPBL ChIP signal at TetO-convergent CTCF sites in TACL-ON cells. Average ChIP signals are normalized to the average signal at the convergent shared peaks_located inside the collapsed domains **(G)** Aggregate FLAG ChIP signal at TetO-convergent CTCF sites in TACL-ON cells. Average ChIP signals are normalized to the average signal at the convergent shared peaks_located inside the collapsed domains **(H)** Aggregate 4C signal plots of TACL-OFF and TACL-ON, STAG2-depleted cells within the STAG2-depleted collapsed domain. Average ChIP signals are normalized to the average signal at the convergent shared peaks_located inside the collapsed domains **(I)** Aggregate NIPBL ChIP signal at TetO-convergent CTCF sites in TACL-ON, STAG2-depleted cells. **(J)** Aggregate FLAG ChIP signal at TetO-convergent CTCF sites in TACL-ON, STAG2-depleted cells.

### Also intradomain CTCF sites loop to the TetO boundary

Next, we then investigated how each category of STAG-defined CTCF peaks contributed to chromatin loops formed with TetO. For this we performed an aggregate analysis of loops formed with TetO: we used the 4C-seq chromatin contact profiles of TetO to select the 4C signals -/+50 kb centered around each of the flanking CTCF sites and we aggregated these signals per category of CTCF sites. We also considered the orientation of their CTCF binding motif to split the data, such that for each category of CTCF sites we could analyze their looping to TetO when oriented towards (TetO-convergent) or away (TetO-divergent) from TetO (Fig. 5D). This aggregate TetO-anchored loop analysis showed that in TACL-OFF cells, as expected, none of the categories of CTCF sites showed preferential contacts with TetO. In TACL-ON cells however, the TetO-convergently oriented CTCF sites of all three categories engaged in looping contacts with TetO. This was most prominently seen for category I and II sites, but also the category III sites showed a higher preference to contact TetO than their surrounding sequences (Fig. 5E). Moreover, even in a TetO-divergent orientation, category I and II sites had contact specificity with TetO in TACL-ON cells (Fig. S5F). Thus, analysis of a large number of CTCF sites for their collective capacity to form loops with an exceptionally active chromatin loop extrusion anchor showed that also intradomain CTCF sites often form temporarily stabilized loops with the anchor. The CTCF sites that looped to TetO accumulated NIPBL (Fig. 5F, fig. S5G) and FLAG-tagged MAU2 (Fig. 5G, fig. S5H), with NIPBL accumulating on both sides of the TetO-divergent CTCF sites (Fig. S5G). We propose the latter is evidence for loop collisions, which would imply that loop extrusion is halted *in vivo* when cohesin encounters another cohesin complex bound to CTCF. The fact that we also find (weak) aggregate looping between TetO and such ‘wrongly’ oriented CTCF sites may well be the consequence of such loop collision events.

We then asked whether the capacity of the three categories of TetO-convergent CTCF sites to loop to TetO altered upon depletion of STAG2. The category I and II sites (the latter now recruiting cohesin^STAG1^instead of cohesin^STAG2^), continued to form specific loops with TetO (Fig. 5H) and accumulate NIPBL (Fig. 5I) and FLAG-tagged MAU2 (Fig. 5J). The category III sites, where cohesin^STAG1^ failed to replace cohesin^STAG2^, lost their ability to loop to TetO (Fig. 5H). They also no longer recruited NIPBL (Fig. 5I) or MAU2 (Fig. 5J). Thus, cohesin^STAG1^ was able to replace cohesin^STAG2^ in engaging some, but not all, intradomain CTCF sites in a looping network. Possibly this inability to involve all CTCF sites in looping events explains why intradomain interactions collapse in the absence of cohesin^STAG2^.

### High cohesin loop extrusion activity hinders transcription

We then investigated whether TACL-induced activated loop extrusion had an impact on the transcription of genes surrounding TetO. We measured and compared nascent transcription in TACL-ON and TACL-OFF cells, as well as in Cherry cells treated with and without Dox (two hours). Nineteen active genes had a TetO platform integrated in their gene body (often seen with PiggyBac insertions^51^): when TetR-MAU2 or TetR-mCherry was released from their gene body (with Dox), expression of these genes was upregulated (Fig. 6A). The remaining genes located within the TACL domains (i.e. the genes not having TetO integrated in their gene body) were categorized based on their distance to TetO. None of these categories of genes responded to TetR-mCherry release from TetO. However, the genes that were located within 250 Kb from TetO showed a collective increase in transcription in TACL cells when we stopped targeted loop extrusion by Dox addition (Fig. 6A, B). This finding suggested that transcription of genes by the RNA polymerase II (RNAPII) machinery is hindered by increasing numbers of bypassing (or halting) extruding cohesin complexes.

**Figure 6.**
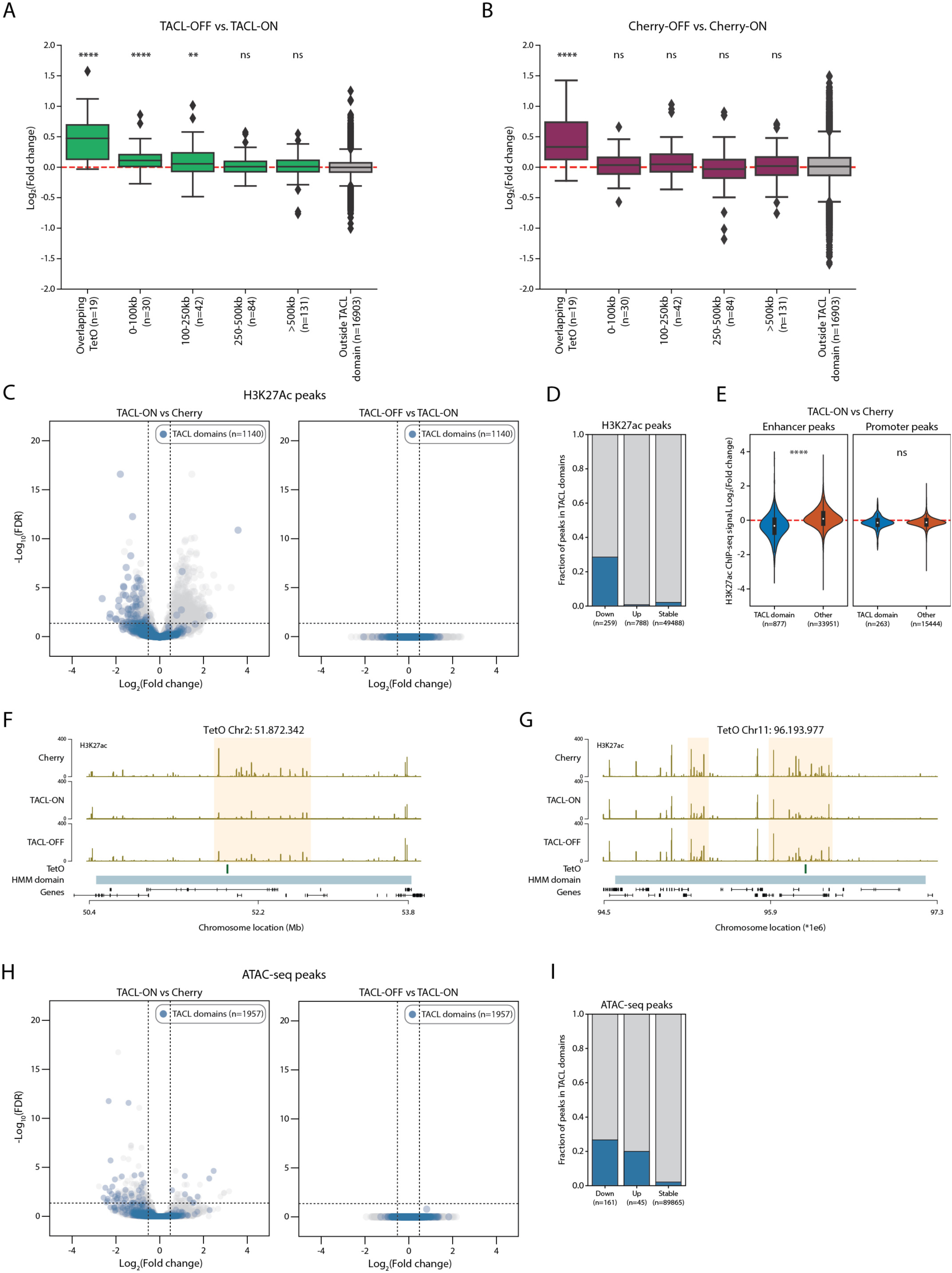
TACL-induced cohesin loop extrusion alters transcription and histone modification. **(A-B)** Relative gene expression changes in TACL-OFF vs TACL-ON (A), or Cherry-OFF vs Cherry-ON cells (doxycycline treated) (B). Y-axis are plotted as log2 fold change between the conditions. Genes within the TACL domain are divided into different groups based on their distances to the TetO sites. 19 genes are overlapping with TetO within their gene bodies. Numbers in the bracket for each group of genes indicates the number of genes in this group. Each group of genes are compared to the control group (outside TACL domain). **** p value < 0.0001; ** p value <0.01 for Mann-Whitney U-test. ‘ns’ indicates not significant. **(C)** Volcano plots depicting differentially-bound H3K27Ac peaks in TACL-ON versus Cherry cells or TACL-OFF verus TACL-ON cells. 1140 peaks are identified within the TACL domain. Note there are more H3K27Ac peaks in TACL domain with decreased levels in TACL-ON compared to Cherry cells. **(D)** Percentages of differentially-bound H3K27Ac peaks in TACL domain. **(E)** Violin plots showing H3K27Ac peaks at enhancer and promoter sites in TACL-ON and Cherry cells. Y-axis are plotted as log2 fold change between the conditions. H3K27Ac peaks are divided into either inside or outside TACL domain. **** p value < 0.0001; ‘ns’ indicates not significant. **(F-G)** H3K27Ac ChIP-seq tracks of two example TetO sites. Note the decreased H3K27Ac levels in TACL-ON cells within the orange square. **(H)** Volcano plots depicting differential ATAC-seq peaks in TACL-ON versus Cherry cells or TACL-OFF verus TACL-ON cells. 1957 peaks are identified within the TACL domain. Note there are more ATAC-seq peaks in TACL domain with decreased levels in TACL-ON compared to Cherry cells. **(I)** Percentages of differential ATAC-seq peaks in TACL domain.

### Cohesin loop extrusion can impact on chromatin accessibility and histone modifications

Finally, we employed TACL to investigate whether locally induced chromatin loop extrusion alter local chromatin accessibility and the epigenetic landscapes. For this, we performed ATAC-seq and ChIP-seq for the active histone marks H3K27ac and H3K4me3, in TACL-On and Cherry control cells. When analyzing H3K27ac signals, we identified 259 lost and 788 gained H3K27ac peaks (out of >50,000) in TACL-On compared to Cherry cells. The sites with gained H3K27ac were not enriched inside TACL domains, but sites with lower levels of H3K27ac were highly enriched in the TACL domains (Fig. 6C, D). This local loss of H3K27ac signal occurred at putative enhancers but not at promoters within TACL domains (Fig. 6E and S6A, B). Promoters also did not alter their H3K4me3 levels because of TACL (Fig. S6C, D). Switching the TACL system off by addition of doxycycline for one hour did not restore the H3K27ac levels (Fig. 6F, G and Fig. S6E). A similarly subtle but significant effect was also seen when analyzing chromatin accessibility by ATAC-seq. Out of ∼90,000 identified accessible sites, only 45 and 161 sites showed a significant gain and loss of accessibility, respectively, in TACL versus Cherry cells. Both categories were highly enriched in TACL domains however, and were not reversible within an hour of Dox treatment (Fig. 6H, I). Our data therefore suggested that prolonged exposure to activated cohesin loop extrusion can have impact on the accessibility and the epigenetic landscape of chromatin, causing mild alterations in H3K27ac levels and the accessibility of potential regulatory sites.

## Discussion

TACL, the system that we generated for targeted recruitment of extruding cohesin complexes, enabled highly sensitive measurements of the impact of cohesin loop extrusion in living cells, for several reasons. By working in haploid eHAP1 cells, we circumvented the problem of simultaneously measuring the status of non-distinguishable untargeted chromosome copies, which could obscure the detection of TACL-induced effects. By selecting a cell line with 27 TetO platforms integrated across nearly all chromosomes, we could go beyond anecdotal single locus observations and perform much more systematic analyses. The TACL domains together spanned ∼68 Mb, or about 2% of the genome. The remainder of the genome served as reference sequences in all our experiments, providing ample internal controls for accurate and sensitive detection of TACL-induced effects. The doxycycline sensitive TetO-TetR protein recruitment system enabled comparing the consequences of activating and not activating local loop extrusion in the same cells. A disadvantage of the TACL system was that we remained ignorant about its absolute activity: we do not know how many loop extrusion events per time unit were initiated from all sites collectively, let alone from each individual site in each given direction. Such measurements are also lacking for naturally initiated loop extrusion events though. Given the strength of TACL-induced chromatin loops and the exceptionally high levels of MAU2 and NIPBL that TACL deposited on flanking CTCF sites, we expect that TACL fired extruding cohesin complexes with very high rates, possibly to also create nested loops in the surroundings of TetO sites.

With TACL we could demonstrate that activated cohesin loop extrusion negatively influenced gene expression and also reduced H3K27ac levels of sites in the TACL domains. Although the effect sizes of these changes were small, this demonstrated that the extruding cohesin machinery can influence the epigenetic make-up of chromatin and the activity of genes that it passes. We currently have no evidence that there is a causal relationship between the two observations. It seems intuitive to interpret the decrease in H3K27ac levels as an indication for reduced local enhancer activity, but we don’t expect that all responding genes need enhancers for their expression. Rather, we believe that gene transcription may be hampered from encounters between the RNA polymerase transcription machinery and the cohesin loop extrusion machinery, which both traverse along the chromatin template. We currently do not know how frequent these encounters are, which will not only depend on the density of loop extruding cohesin complexes on chromatin, but also on gene sizes and their (unknown) frequencies of transcriptional bursts. We recently already reported that cohesin can hamper gene transcription. This was based on the more indirect observation that an enhancer, shown to recruit cohesin, required cohesin for long-range gene activation but was antagonized by cohesin for short-range gene activation^36^. A direct link between cohesin-mediated loop extrusion and the accessibility and epigenetic make-up of chromatin is interesting: it suggests that cohesin loop extrusion may fine-tune the activity of chromatin.

Why sites lose K27ac when exposed to high cohesin loop extrusion activity remains to be investigated, but it is possible that steric hindrance with chromatin entrapped by cohesin being temporarily less accessible to *trans*-acting factors supporting H3K27 acetylation.

An unexpected observation was the TACL-induced, cohesin-dependent accumulation of NIPBL and MAU2 at flanking CTCF sites. This was found predominantly at convergent CTCF sites, but also at divergent CTCF sites, where cohesin appeared to accumulate at both sides of the CTCF binding site. We propose the latter was the consequence of oppositely extruding cohesin molecules colliding and pausing at these sites. This may also explain why, under these activated loop extrusion conditions, these divergent sites were found looping to TetO. NIPBL-MAU2 recruitment to CTCF sites was not a peculiarity of the TACL system, as we also observed this happening elsewhere in the genome, both in TACL and in non-TACL (Cherry) cells, and in datasets published by others^12^. It was best appreciable in TACL cells though, also outside the TACL domains. This could be because of the 1.8-fold elevated nuclear MAU2 levels present in TACL cells. Alternatively, TetR-MAU2 molecules may be better cross-linkable to DNA than normal MAU2, facilitating their detection by chromatin immunoprecipitation. Because we found a strong correlation between NIPBL-MAU2 association to CTCF sites and their looping to TetO, we hypothesize that cohesin-mediated CTCF-CTCF loop formation and stabilization requires continued active support from associated NIPBL-MAU2.

*In vitro*, an exchange of NIPBL molecule binding to extruding cohesin was reported to be required for continued loop extrusion^19,20^, but we found no evidence for MAU2 molecules replacing each other on cohesin extruding from TetO in TACL cells. This may suggest different binding dynamics of NIPBL and MAU2, the two auxiliary factors. It may also be that in TACL cells the TetR-MAU2 protein concentrations locally around TetO were too high for MAU2-V5 to effectively compete in the exchange. Alternatively, we studied anchored cohesin loop extrusion (acting on chromatinized DNA) whereas the *in vitro* work studied unanchored cohesin loop extrusion (acting on naked DNA). Possibly, anchored cohesin remains stably associated with NIPBL-MAU2 for the reeling in of flanking sequences and the formation of stabilized chromatin loops with other anchors.

In our TACL system, cohesin^STAG2^ appeared to be the dominant protein complex recruited to TetO. Cohesin^STAG2^ was previously demonstrated to be required for interactions between enhancers and promoters inside TADs, a function that could not be taken over by cohesin^STAG1^ when STAG2 was depleted^11,12^. Our data confirmed that only cohesin^STAG2^ supported chromatin contacts within the TACL-induced loop domains, and provided possible explanations for this. We found that most, if not all, CTCF-bound DNA sites have the capacity to halt cohesin^STAG2^, and to form temporarily stabilized chromatin loops with other loop anchors. We expect that such transiently stabilized intradomain CTCF-CTCF interactions are constantly formed everywhere in the genome. Without TACL tremendously boosting loop extrusion from defined anchors though, and without the aggregate anchored loop analysis used here, they will likely remain undetected: each of these individual loops will be formed too rarely among the majority of un-looped alleles to be discovered in cell population-based Hi-C or 4C analyses, or to be appreciable in single-cell Hi-C analysis. A complex network in which all CTCF sites dynamically participate in intradomain CTCF-CTCF loops, we propose, may well be essential to support intradomain contacts. In STAG2-depleted cells, cohesin^STAG1^ physically and functionally replaced cohesin^STAG2^ at some, but certainly not at all intradomain CTCF sites. Without cohesin^STAG2^, these ‘STAG2-only’ sites (category III; the weaker CTCF-binding sites) lost their capacity to recruit NIPBL-MAU2 and lost their ability to engage in stabilized contacts with other loop anchors. This inability of cohesin^STAG1^ to engage all intradomain CTCF sites in a dynamic network of CTCF-anchored loops may well explain the collapse of intradomain contacts seen upon STAG2 depletion. It is tempting to speculate that this is also a reason why cohesin^STAG1^ associates more stably with chromatin than cohesin^STAG2^. Assuming that cohesin^STAG2^ is released from chromatin predominantly when stalled at these intradomain CTCF sites (where also WAPL and PDS5A are found^29^, and where cohesin strongly accumulates in the absence of WAPL^22^), cohesin^STAG2^ would be more quickly released because it pauses at more CTCF sites than cohesin^STAG1^. CTCF, in such scenario, would then have two functions: at binding sites where it is optimally oriented for its N-terminus to face and stably interact with the conserved essential surface (CES) of incoming cohesin^24^ it will prevent WAPL from binding to the CES and releasing cohesin from chromatin^22,52,53^. At other DNA binding sites, perhaps because of its suboptimal positioning, it may halt but not sufficiently stably associate with cohesin, facilitating WAPL to bind to the CES and release cohesin from chromatin.

With TACL now enabling controlled targeted activation of cohesin loop extrusion trajectories, we expect new avenues are opened to study in detail and possibly even visualize individual loop extrusion trajectories, the consequences of their encounters with each other and with the transcription machinery, their impact on gene regulation by distant enhancers and their interplay with all different types of epigenetic chromatin landscapes *in vivo*.

## Methods

### Cell culture

eHAP1 cells were cultured in Iscove’s Modified Dulbecco’s Medium (IMDM) medium supplemented with Glutamax (Thermofisher), 25mM Hepes, 10% FBS, and 1% Penicillin-Streptomycin following standard procedures. Cells were routinely checked and sorted for haploidy. 293TX cells were cultured in Dulbecco’s Modified Eagle Medium (DMEM) medium supplemented with 10% FBS and 1% Penicillin-Streptomycin.

### Antibodies

Anti-SMC1 (A300-055A, Bethyl), anti-SMC3 (A300-060A, Bethyl), anti-RAD21 (05-908, Merck), anti-NIPBL (A301-779A, Bethyl), anti-FLAG (F1804, Merck), anti-SCC4/MAU2 (ab183033, Abcam), anti-GAPDH (sc-32233, Santa Cruz), anti-STAG1 (A302-579A, Bethyl), anti-STAG2 (A300-159A, Bethyl), anti-CTCF (ab128873, Abcam), anti-H3K4me3 (39060, Active motif), anti-H3K27ac (39133, Active motif), anti-V5 (R960-25, Thermo fisher), anti-WAPL (sc-365189, Santa Cruz), anti-PDS5A (A300-089A, Bethyl).

### Plasmid construction

The plasmids expressing TetR-FLAG-MAU2 and TetR-FLAG-mCherry cassette was cloned into a Lentivirus backbone under the control of EF1 promoter. TetR, FLAG, and MAU2/mCherry sequences were PCR amplified with 20bp overhang for In-Fusion cloning. The final expression cassette is composed of EF1-TetR-FLAG-MAU2/mCherry-P2A-Puromycin. To construct the V5-MAU2 plasmid, TetR-FLAG sequence from the TetR-FLAG-MAU2 construct was removed, and a V5 tag was inserted instead. To enable simultaneous expression of the two MAU2 constructs, the antibiotic selection marker was replaced by blasticidin instead of puromycin. To insert the auxin inducible degron (AID2) tag into the endogenous gene, a sgRNA targeting the ORF of the gene was cloned into a vector containing SpCas9-T2A-BFP (Table S1). To construct the donor template for AID2 tag insertion, a cassette containing AID2-GFP was cloned between two homology arms of about 1kb surrounding the sgRNA cut site.

### Generation of cell lines containing the TetO platforms

The plasmids bearing the TetO platforms and the piggybac transposase were originally obtained from Luca Giorgetti^39^. Nanopore validation 48x repeats Briefly, eHap1 cells were trypsinized and resuspended in serum-free IMDM medium. A vector containing the piggybac transposase (pBroad3_hyPBase_IRES_tagRFPt) were mixed with a piggybac donor vector bearing 30x TetO binding sites and Polyethylenimine (PEI, polysciences) in serum-free IMDM. The DNA mix was incubated at room temperature for 10min, after which the cells and the DNA mix were incubated together for another 10min. The cells were then plated in a 6-well plate. After 24h, the medium was refreshed. 48-72h after the transfection, the cells were sorted for RFP signal, expressing the transposase. Sorted cells were plated in 15cm dish and cultured for at least 14 days. Colonies were picked and sub-cultured in 96-well plates. To genotype the clones with a sufficient number of integration sites, cells were lysed in DirectPCR lysis reagent (Viagen). Lyastes were subsequently assessed by running qPCR with primers annealing to the transposon sequences. A primer targeting a part of human FSIP2 gene was used as the reference among different clones. An estimation of the number of integration sites was calculated as: 2^-(Ct_TetO primer – Ct_reference). The exact number of integrations sites was validated by 4C-seq. Integration site mapping is missing.

### Lentivirus production and transduction

4×10^6^ 293TX cells were plated in 10cm dish 24h prior virus production. Lentiviral vectors were co-transfected with pVSV-G, pMDL RRE, and pRSV-REV in serum-free DMEM with Polyethylenimine (PEI, polysciences). The medium was refreshed 18h after transfection. The medium containing the virus particles were harvested 48h after transfection by passing through a 0.45μm filter. For transduction, eHap1 cells were plated in a 6-well plate 24h before transduction. The transduction was performed by adding the virus particle directly onto the cells supplemented with 6μg/mL polybrene (Merck). The cells were refreshed 24h after transduction and the antibiotics (puromycin/blasticidin) were added 48h after transduction. Cells were selected with antibiotics until the cells in the control plate (without transduction) were completely dead.

### Western blot

Cells were washed in PBS and lysed in RIPA buffer with protease inhibitor (Roche) on ice for 15min. The cell lysate was further disrupted by sonication with Bioruptor Pico (Diagnode). The cell lysate was cleared by spinning at 1000xg for 5min. The supernatant was incubated with Laemmli buffer and boiled for 10min. The sample was then loaded on a 4–15% Mini-PROTEAN® TGX™ Precast Protein Gel (Biorad) and ran at 100V for 90min. Proteins were transferred onto a nitrocellulose or PVDF membrane and incubated with the primary antibody overnight at 4°C. The membrane was then washed in PBS-0.25% Tween and incubated with the secondary antibody at room temperature for 1h. Finally, the membrane was incubated with SuperSignal™ West Pico PLUS Chemiluminescent Substrate (Thermofisher) for 1min before visualized on ImageQuant 800 imager (Amersham).

### Nuclear and cytoplasmic fractionation

Briefly, 3×106 cells were collected by trypsinization. Cells were washed with PBS and cell pellet was resuspended in 100μL of Cytoplasmic extraction buffer (10mM HEPES, 60 mM KCl, 1 mM EDTA, 0.075% (v/v) NP-40, 1mM DTT and 1 mM PMSF, final pH 7.6) and incubated on ice for 3min. Suspension was spun at 1500rpm for 4min and the supernatant was kept as cytoplasmic fraction. The pellet was washed once with Cytoplasmic extraction buffer. Cells were pelleted at 1500rpm for 4min and resuspended in 50μL of Nuclear extraction buffer (20 mM Tris Cl, 420 mM NaCl, 1.5 mM MgCl2, 0.2 mM EDTA, 1 mM PMSF and 25% (v/v) glycerol, final pH 8.0). Adjust the salt concentration to 400mM of NaCl and add an additional pellet volume of Nuclear extraction buffer. Pellet was vortexed and incubated on ice for 10min. The suspension was spun at max speed for 10min and supernatant was kept as nuclear fraction.

### Chromatin immunoprecipitation (ChIP)

100 million cells were crosslinked with 1% formaldehyde for 10min. Cells were subsequently quenched with 125mM glycine for 10min and washed twice with cold PBS. Cells were scraped from culture dishes and cell pellets were subsequently lysed in LB1 buffer (50mM Hepes, 140mM NaCl, 1mM EDTA, 10% glycerol, 0.5% NP40, 0.25% Triton X-100), washed in LB2 buffer (10mM Tris, 200mM NaCl, 1mM EDTA, 0.5mM EGTA), and resuspended in LB3 buffer (10mM Tris, 100mM NaCl, 1mM EDTA, 0.5mM EGTA, 0.1% sodium deoxycholate, 0.5% N-lauroylsarcosine) prior sonication. Chromatin was sonicated using Bioruptor Pico (Diagnode) with a setting of 30s on, 30s off for 8 cycles. Fragmented chromatin was then incubated with 6ug of antibodies pre-coupled to Dynabeads protein G beads (Thermofisher) overnight at 4°C. Beads-bound chromatin was then washed 10x with RIPA buffer (50mM Hepes, 500mM LiCl, 1mM EDTA, 1% NP40, 0.7% sodium deoxycholate), once with TBS buffer, and decrosslinked in elution buffer (50mM Tris, 10mM EDTA, 1% SDS) at 65°C for 18h. Eluted DNA was then treated with protease K and RNAse A, and subsequently purified with phenol/chloroform/isoamyl alcohol 25:24:1. Purified DNA was either assessed with qPCR or continued with ChIP-seq NGS sequencing library preparation. Sequencing libraries was constructed using NEBnext Ultra II DNA library prep kit (NEB) following the manufacture’s protocol. Briefly, DNA was end-repaired and poly-A tailed, ligated to NEBnext adapters, and digested with USER enzyme. Annealed libraries were then purified with AMPure XP beads (Beckman Coulter) and PCR amplified with indexing primers for 4-12 cycles. Sequencing libraries were checked with Bioanalyzer HS DNA chip (Agilent) and sequenced on the Illumina Nextseq 500 (single end reads, 75bp) and Nextseq 2000 platforms (pair end reads, 50bp).

### 4C-seq

4C templates preparation was performed as described^54,55^. In brief, ten million cells per sample were crosslinked with 2% formaldehyde, followed by quenching by glycine at final concentration of 0.125 M. Four-cutter restriction enzyme MboI (New England BioLabs) was used for in situ digestion (300U/10 million cells). Digested DNA fragments were ligated, reverse-crosslinked and subsequently purified through isopropanol and magnetic beads (Macherey-Nagel NucleoMag PCR Beads). Four-cutter restriction enzyme Csp6I (CviQI, ThermoFisher ER0211, 50U/sample) was used for template trimming. Re-ligated and purified 4C templates were further proceed through *in vitro* Cas9 digestion as described below.

### *In vitro* Cas9 digestion of 4C templates

To prevent PCR amplification and sequencing of TetO repeats due to tandem ligation of two or more TetO DpnII fragments in a given 4C circle, an *in vitro* digestion of 4C templates was performed as described in^56^ with the following modifications: two sgRNA were used to target Cas9 into the TetO repeats between viewpoint primers; pre-incubation of Cas9 protein and sgRNA template at room temperature. In brief, two sgRNA templates were obtained using the Megashortscript T7 transcription kit (Invitrogen) followed by 4x AMPure XP (Agencourt) purification. Purified Cas9 protein (generated by Hubrecht protein facility) was pre-incubated with the sgRNAs for 30min at room temperature. 4C templates were subsequently added to the pre-incubated Cas9/sgRNA complexed for overnight digestion at 37°C. Cas9 protein was inactivated by incubating at 70 °C for 5 min. Resulting products were purified with 1x AMPure XP and used as PCR template for TetO dedicated 4C.

### Nascent RNA sequencing (BrU-seq)

BrU-seq was performed as described^57^. Cultured cells were incubated with 2mM Bromouridine (BrU, Merck) for 10min and subsequently lysed in TRIzol reagent (Thermofisher). RNA was isolated following the manufacture’s protocol. Briefly, lysed cells were mixed with chloroform and centrifuged for 15min. The aqueous phase was transferred to a new tube and mixed with isopropanol. After centrifugation, RNA pellet was washed once with 70% ethanol and dissolved in DEPC water. To capture BrU-labelled nascent RNA, 6ug anti-BrdU antibodies (BD biosciences) pre-coupled with Dynabeads protein G beads (Thermofisher) were incubated with the total RNA for 1h at room temperature. Beads were then washed 3x with PBS/0.1% Tween-20/RNaseOUT. To purify the beads-bound RNA, TRIzol reagent was directly added to the beads and RNA was purified as described above. NGS sequencing libraries were generated using NEBnext Ultra II directional RNA library prep kit (NEB) following manufacture’s protocol. Briefly, RNA was fragmented to about 200bp in size. First strand and second strand cDNA were synthesized. Double strand cDNA was then end repaired, poly-A tailed, ligated to NEBnext adapters, and digested with USER enzyme. Annealed libraries were then purified with AMPure XP beads (Beckman Coulter) and PCR amplified with indexing primers for 7 cycles. Sequencing libraries were checked with Bioanalyzer HS DNA chip (Agilent) and sequenced on the Illumina Nextseq 2000 platforms (single end reads, 50bp).

### Hi-C

Hi-C template preparation was performed as described in^25^. In brief, ten million cells per sample were crosslinked with 2% formaldehyde followed by quenching by glycine at final concentration of 0.2 M. Four-cutter restriction enzyme DpnII (New England BioLabs) was used for in situ digestion (400U/ 10 million cells). Digested DNA were repaired with biotin-14–dATP (Life Technologies) in a Klenow end-filling reaction. End-repaired, ligated, and reverse-crosslinked DNA was subsequently purified using isopropanol and magnetic beads (Macherey-Nagel NucleoMag PCR Beads). Purified DNA was sheared to 300-500 bp with Covaris and subsequently size-selected by AMPure XP (Agencourt). Appropriately-sized ligation fragments marked by biotin were pulled down with MyOne Streptavidin C1 DynaBeads (Invitrogen) and prepped for Illumina sequencing.

### ATAC-seq

ATAC-seq was conducted following the Omni-ATAC protocol. In summary, 200,000 cells were lysed using a solution containing 0.1% NP-40, 0.1% Tween-20, and 0.01% digitonin, then incubated with a homemade Tagment DNA Enzyme for 30 minutes at 37 °C. DNA purification was carried out using the QIAGEN MinElute Reaction Cleanup Kit. Library fragments were amplified with Phusion High-Fidelity PCR Master Mix with HF Buffer (Thermo Fisher Scientific, catalog no. F531S) and custom primers featuring unique single or dual indexes. Purification of the libraries was performed using AMPure XP beads (Beckman Coulter, catalog no. A63881), following the manufacturer’s guidelines. The quality of the constructed libraries was assessed using the Agilent Bioanalyzer 2100 with the DNA 7500 kit (catalog no. 5067-1504).

### Generation of auxin-inducible degron cells

To deplete the cohesin factors in cells, we utilized the AID2 system^44^. For RAD21 degron, we generated eHap1 cells stably expressing OsTIR1 (F74G) by transducing the cells with lentivirus containing an expression cassette of OSTIR1-P2A-hygromycin. After antibiotic selection with hygromycin, cells were co-transfected with a vector expressing a sgRNA against RAD21 and SpCas9-T2A-BFP, and the donor template containing AID-GFP flanked by homology arms. GFP-positive cells were analyzed and sorted with flow cytometry. Single cell clones were expanded and used for downstream analysis. For WAPL, PDS5A, STAG2, and CTCF degrons, we first inserted the AID-GFP cassette by co-transfecting the cells with a sgRNA against each gene. Single cell clones were selected and verified by PCR. Verified clones were then transduced with lentivirus containing an expression cassette of OSTIR1-P2A-blasticidin. To deplete the proteins, we used auxin (5-Ph-IAA, BioAcademia) and treated the cells for 2-3h and analyzed the successful depletion with western blot.

## Data analysis

### 4C-seq

4C-seq reads were mapped to the hg38 reference genome and processed using pipe4C^54^ ( https://github.com/deLaatLab/pipe4C). Normalized 4C coverage was calculated separately for each TetO integration site using R (r-project.org). Counts at non-blind fragments within a 20 MB region (10 MB upstream and downstream of the viewpoint) were adjusted to 1 million mapped reads after exclusion of the two highest-count fragments. Count data was smoothed using a running mean with a window size of 21 fragments using the R package caTools v 1.18.2.

#### Aggregate 4C analysis

In 3C-based assays, ligation frequencies are typically highest near the viewpoint (<100 kb) and decrease as the distance from the viewpoint increases. To minimize the high background ligation frequencies close to the viewpoint, only peaks located at least 100 kb away from the TetO integration sites were included in the aggregate 4C analysis. These peaks were resized to 100 kb, divided into 2 kb bins, and the average normalized 4C signal was calculated for each bin.

#### TACL domains annotation

To systematically annotate the TACL domains induced by the recruitment of cohesin to the TetO platforms, we developed a Hidden Markov Model (HMM). The HMM was implemented using the Python package hmmlearn (https://github.com/hmmlearn/hmmlearn). We created an HMM with the states ‘TACL_domain’ and ‘no_change’. The normalised 4C-seq signals for TACL-ON and Cherry conditions were binarised into two observations: “*TACL_domain*” (4C-seq signal difference between TACL-ON and Cherry greater than 25) and “*no_change*” (4C-seq signal difference less than or equal to 25). The emission probabilities were estimated using manually defined TACL domains. The probability of the “*TACL_domain*” state was calculated as the fraction of restriction fragments with 4C-seq signal greater than 25 in the manually defined TACL domains and set to 0.6. The probability of the “*no_change*” state was calculated as the fraction of restriction fragments with 4C-seq signal less than or equal to 25 in the flanking regions of the manually defined TACL domains and was set to 0.98. The transition probability was set to 10^-6^.

The estimated “*TACL_domain*” and “*no_change*” states were then subjected to several additional filters. First, restriction fragments belonging to stretches of more than 20 consecutive “*TACL_domain*” states were retained. Second, restriction fragments with consecutive “*TACL_domain*” states within 100 kb of each other were merged. Third, merged regions containing at least 40 restriction fragments were retained and further merged within 1.5 Mb of each other to draft TACL domains. Finally, if TetO was outside of the drafted TACL domain, the closest domain segment on the other side of the domain with respect to the TetO location was added to obtain TACL domains.

HMM model with the same parameters was used to annotate TACL domains in the CTCF-AID, WAPL-AID, STAG2-AID, and PDS5A-AID lines by comparing the difference between IAA and Dox treatments. For the filtering steps, the distance for considering restriction fragments with consecutive “*TACL_domain*” states was set to 200 kb and the distance for drafting TACL domains from restriction fragments was set to 2.5 Mb.

### ChIP-seq

Publicly available HAP1 H3K4me1 data was included [REF}.

ChIP-seq reads were mapped to the hg38 reference genome and processed using the 4DN ChIP-seq pipeline (https://github.com/4dn-dcic/chip-seq-pipeline2). P-val signal bigwigs were used for all heatmaps and example plots. For WT, T-MAU2, T-MAU2 treated with doxycycline or T-mCherry cells, the p-val signals were normalized based on the average p-val signal for all CTCF peaks in TACL-ON (for CTCF, RAD21, SMC1, SMC3, STAG1, STAG2, WAPL, PDS5A), FLAG peaks in TACL-ON (for FLAG, MAU2, NIPBL, V5), H3K27ac peaks (H3K27ac) or H3K4me3 peaks (H3K4me3) located outside the TACl domains and further than 3MB from the TetO integration sites. Briefly,ChIPseq peaks were filtered for a signalValue which represented clear peaks by visual inspection (CTCF: 35, FLAG-MAU2: 35) and for overlapping peaks, such that for overlapping peaks the peak with the highest signalValue was kept. Filtered peaks were resized to 10 bp and the signal was calculated using the GenomicRanges and rtracklayer R/Bioconductor^58,59^. The average signal was used as the scaling factor. For degron lines p-val signals were normalized based on the average signal of the regions flanking the filtered peaks. Briefly, peaks were filtered for signalValue and for overlapping peaks as described above. Next, peaks were resized to 5kb and the signals of the outer 1kb regions (2.5kb-1.5kb upstream and downstream of the peak center) were calculated. The average signal of the outer 1kb regions was used as the scaling factor. For heatmaps, the signal coverage was calculated per 10 bp bin as described above and normalized using the previously determined scaling factor. For the average ChIP signal plot the average signal for each 10 bp bin was calculated.TetO enrichment ChIP-seq reads were mapped to the hg38 human reference genome assembly with added minimal PiggyBac TetO sequence (Table S2) using bowtie2 v2.5.2 (PMID: 22388286). Alignments with a mapping quality (MQ) score of ≥1, either to the PiggyBac TetO sequence or elsewhere in the genome, were quantified using FeatureCounts v2.0.6 (PMID: 24227677). Enrichment levels were determined by comparing the coverage to the average coverage from all input control experiments.

#### Differential FLAG peaks

FLAG ChIP-seq reads were mapped as SE reads to the hg38 human reference genome assembly with added TetO sequence using bowtie2 v2.5.2 (PMID: 22388286). Uniquely mapped reads with MAPQ >= 15 were selected using SAMtools v1.15 (PMID: 19505943). Duplicate reads were filtered out using the Picard v2.25.6 and v3.1.1 “MarkDuplicates” function (https://broadinstitute.github.io/picard/). Coverage at FLAG peaks was counted using FeatureCounts v2.0.6 (REF) and normalized using DESeq2 v1.38.3 (REF), Differential FLAG peaks were defined as peaks with M>1 and dFLAG <- merged_GR[merged_GR$M > 1 & (merged_GR$on_avg + 1) > 2^4.5, ]

#### CTCF motif analysis

The presence and orientation of CTCF motifs below each CTCF peak were identified using FIMO v5.3.0^60^ with motif MA0139.1^61^ and max-stored-scores 50000000. CTCF peaks for which all identified motifs were located on the plus strand were classified as forward CTCF peaks, while peaks for which all identified motifs were located on the minus strand were classified as reverse CTCF peaks. Forward CTCF motifs located upstream of TetO sites and reverse CTCF motifs located downstream of CTCF were classified as convergent CTCF binding sites. Reverse CTCF motifs located upstream of TetO sites and forward CTCF motifs located downstream of CTCFwere classified as divergent CTCF binding sites.

### Classification of STAG2 Peaks

Genome-wide shared STAG1/2 sites are defined as STAG2 peaks located outside TACL domains and situated at least 3 MB away from any TetO integration site in TACL-ON, and that overlap with STAG1 peaks in both TACL-ON and dSTAG2 TACL-ON. Category I (Shared STAG1/2 sites), Category II (Conditional STAG1 sites) and Category III (STAG2-only sites) STAG2 sites are STAG2 peaks in TACL-ON that overlap with CTCF peaks and which are located within the TACL domains. Category I (Shared STAG1/2 sites) overlap with STAG1 in dSTAG2 TACL-ON and also overlap with STAG1 peaks in TACL-ON,or have a STAG1 coverage ≥ the 10th percentile of the genome-wide shared STAG1/2 sites. Category II (Conditional STAG1 sites) overlap with STAG1 in dSTAG2 TACL-ON but do not overlap with STAG1 peaks in TACL-ON, with STAG1 coverage < the 10th percentile of genome-wide shared STAG1/2 sites. Category III (STAG2-only sites) do not overlap with STAG1 peaks in either TACL-ON or dSTAG2 TACL-ON.

### Anlaysis of ATAC-Seq and ChIP-seq for histone modifications

#### Data processing

ATAC-seq reads were mapped to the hg38 human reference genome assembly using bwa mem v0.7.17-r1188 (PMID: <u>19451168</u>). ChIP-seq reads were mapped to the hg38 human reference genome assembly with added TetO sequence using bowtie2 v2.5.2 (PMID: 22388286). Uniquely mapped reads in proper read pairs (-f 2) with MAPQ > 10 and MAPQ >= 15 were selected using SAMtools v1.15 (PMID: 19505943) for ATAC-seq and ChIP-seq data, respectively. Duplicate reads were filtered out using the Picard v2.25.6 and v3.1.1 “MarkDuplicates” function (https://broadinstitute.github.io/picard/) for ATAC-seq and ChIP-seq data, respectively. Bigwig coverage tracks were generated using the “bamCoverage” function from the deepTools v3.4.2 (PMID: 27079975) with the “--effectiveGenomeSize” parameter set to 2913022398 and “--normalizeUsing” parameter set to RPGC.

#### Peak calling

Peaks were called using MACS2 v2.2.6 (PMID: <u>18798982</u>) for pooled data and replicates in a narrowPeak mode, with mappable genome size set to hs, q-value cutoff of 0.05, “--keep-dup” parameter set to all, and “--nomodel” parameter. The consensus peak list was obtained by overlapping the peaks called for pooled data with peaks from replicates. Only the peaks from canonical chromosomes outside of the blacklist regions (PMID: <u>31249361</u>) that had an overlap of at least 50% with peaks from both replicates were retained.

#### Peak analysis

ATAC-seq and H3K27ac peaks from the TACL-ON, TACL-OFF and Cherry conditions were pooled into one set for differential occupancy analysis. Peak counts were obtained using the “intersect” function from BEDTools v2.27.1 (PMID: <u>20110278</u>) with “-c -wa” parameters.

Differential ATAC-seq and H3K27ac peaks were identified using the DESeq2 v1.30.1 (PMID: 25516281). The “nbinomWaldTest” function with default parameters was used to test contrasts. Peaks with FDR < 0.05 and absolute log2-fold-change > 0.5 were considered significant. For downstream analyses, peak overlap was performed using bioframe v0.3.0. H3K27ac peaks that overlapped with H3K4me3 peaks were classified as promoter peaks, and non-overlapping peaks were classified as enhancer peaks.

### Bru-seq

#### Data processing

BrU-seq reads were mapped to the hg38 human reference genome assembly using STAR v2.7.9a (PMID: 23104886) with GENCODE v44 gene annotation. Uniquely mapped reads with MAPQ > 10 were selected and split by strand using SAMtools v1.12 (PMID: 19505943). Forward strand reads were extracted by using -f 16 FLAG, reverse strand reads were extracted by using -F 16 FLAG. Gene counts were obtained using the “htseq-count” function from HTSeq v0.13.5 (PMID: 25260700). Counts were calculated separately for genes from forward and reverse strands with the parameters “--stranded no”, “--nonunique all”, “--order pos”, and “--type gene”.

#### Differential expression analysis

Differentially expressed genes were identified using the DESeq2 v1.30.1 (PMID: 25516281) (Table S3). Low expressed genes were filtered by requiring the samples to have gene counts greater than 10. The “nbinomWaldTest” function with default parameters was used to test contrasts. Genes with FDR < 0.05 and absolute log2-fold-change > 1 were considered significant. For downstream analyses, the genes were overlapped with annotated TACL domains and split into groups depending on their relative distance and position to the TetO platforms using bioframe v0.3.0.

### Hi-C analysis

#### Data processing

Hi-C data was processed using the distiller pipeline from Open2C (https://github.com/open2c/distiller-nf). The reads were mapped to the human reference genome assembly hg38 with bwa mem v0.7.17-r1188 (PMID: <u>19451168</u>) with “-SP” FLAGs. The alignments were parsed and filtered for duplicates using the pairtools v0.3.0 (PMID: <u>38809952</u>). The complex walks in long reads were masked with “--walks-policy” set to mask, the maximal allowed mismatch for reads to be considered as duplicates “max_mismatch_bp” was set to 1, and the mapping quality threshold was set to 30. Filtered reads pairs were aggregated into genomic bins of different sizes using the cooler v0.8.11 (PMID: <u>31290943</u>). The resulting Hi-C matrices were normalised using the iterative correction procedure.

#### Loops and TADs annotation

High-resolution Hi-C data for HAP1 cells (PMID: <u>28475897</u>) at 10 kb resolution was used for loops and TADs annotation. Loops were annotated using chromosight v1.4.1 (PMID: <u>33199682</u>). For loop detection the Pearson correlation threshold was set to 0.4, loop sizes were set between 50 kb and 5 Mb, and parameter --smooth-trend was enabled. TADs were annotated using the insulation score algorithm implemented in the cooltools v0.3.2 diamond-insulation function (PMID: <u>38709825</u>). The window size for insulation score calculations was set to 200 kb. The threshold for the boundary strength filter was calculated using the Li method, implemented in the scikit-image package (PMID: <u>25024921</u>). The bins with boundary strength higher than ∼0.19 were considered as TAD boundary bins. These bins were converted into TADs by continuously joining two neighbouring bins together. The TAD boundary coordinate was then randomly selected from the coordinates of the joined bins with significant insulation score.

#### Aggregate analyses

Average loops, TAD boundaries, and TADs were calculated for 10 kb resolution observed-over-expected Hi-C contact matrices using the loops and TADs annotated as described above. Publicly available HAP1 Hi-C data was included for comparison (PMID: <u>26499245</u>, <u>28475897</u>). Expected contact matrices were obtained using cooltools v0.3.2 function “compute-expected” (PMID: <u>38709825</u>). Average loops were generated using coolpup.py v0.9.5 (PMID: <u>32003791</u>) with “pad” set to 200 and “min-dist” set to 0. Average TAD boundaries were generated using coolpup.py v0.9.5 (PMID: <u>32003791</u>) in “local” mode with “pad” set to 500. Average TADs generated using coolpup.py v0.9.5 (PMID: <u>32003791</u>) in “local” mode with “rescale” option, with “rescale_size” set to 99. The average loop strength was calculated as the mean value of the central 3 by 3 square pixels. The average TAD boundary strength was calculated as the mean value of the average intra-TAD interactions (upper-left and bottom-right quarters) divided by the mean value of average inter-TAD interactions (upper-right and bottom-left quarters). The average TAD density was calculated as the mean value of the central 33 by 33 square pixels.

The aggregate stripes analysis of the TetO integrations was performed using cooltools v0.5.1 (PMID: <u>38709825</u>) and bioframe v0.3.0 (PMID: <u>38402507</u>) for 10 kb resolution observed-over-expected Hi-C contact matrices. The pile-ups of the TetO integrations were created using the cooltools.pileup function with 500 kb regions around the integration coordinates as flanks.

## Acknowledgments

We thank L. Giorgetti for providing the initial TetO constructs; M. Tanenbaum for providing the TetR construct; Q. Liu for assistance with cloning at the early stage of the project; C. Valdes and R. Neijts for valuable discussions; R. van der Weide for help with ChIP-seq analysis; and all members of the de Laat lab for support and advice. This work was funded by the NWO Groot grant (2019.012) and co-financed by Oncode Institute, which is partly funded by the Dutch Cancer Society.

## Author contributions

R.H. and W.d.L. conceived and initiated the project. R.H. cloned the TetR-MAU2 construct. R.H. and Y.H. generated the cell lines. R.H. performed the ChIP-seq and BrU-seq experiments. Y.H. performed the 4C-seq experiments. M.V. performed ATAC-seq, western blot and Hi-C experiments. K.Z. and M.R. helped cloning the degron constructs and generating the cell lines. M.R. performed part of the 4C, ChIP-seq and BrU-seq experiments. Y.H., P.H.L.K., and M.M. performed 4C-seq analysis. I.V. and P.H.L.K. performed integration site mapping and ChIP-seq analysis. M.M. and A.A. performed BrU-seq analysis. M.M. and P.H.L.K performed histone modification and ATAC-seq analysis, and Hi-C analysis with input from E.d.W. R.H. and W.d.L. drafted the manuscript with input from other authors. R.H., Y.H., M.R., and P.H.L.K. edited the figure panels.

## Competing interests

Authors declare that they have no competing interests.

## Data and materials availability

Cell lines, plasmids, and other materials are available upon request. Sequencing data are available at GSE218803. The code for the 4C-seq and Hi-C data analysis has been deposited at Github (https://github.com/magnitov/tacl) and is publicly available as of the date of publication.

---

**Figure S1.**
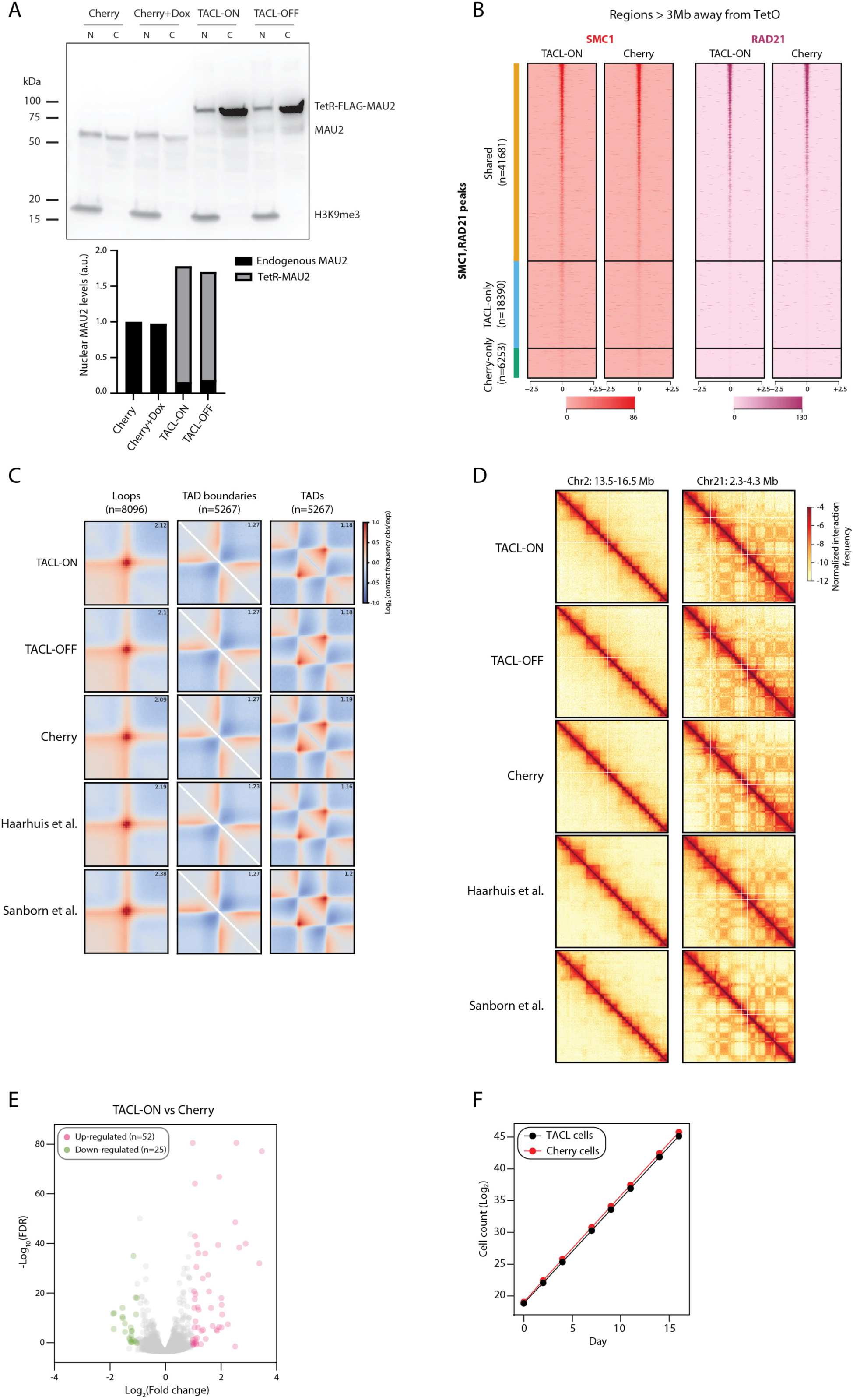
TetR-FLAG-MAU2 functionally replaces endogenous MAU2 and does not cause physiological changes in eHap1 cells. **(A)** Western blot analyzing the nuclear and cytoplasmic fractions of Cherry, Cherry+Dox, TACL-ON, and TACL-OFF cells. Upper panel shows the image from the western blot. H3K9me3 was used as an indicator of the nuclear fraction. ‘N’ represents nuclear and ‘C’ represents cytoplasmic. Lower panel shows the quantification of the nuclear fraction. Intensities of the protein bands were measured and used for calculations. **(B)** Heatmaps of ChIP-seq signals (SMC1 and RAD21 peaks in TACL-OFF, TACL-ON or Cherry cells) that are located outside the TACL-domains and are more than 3Mb away from the nearest TetO integration site in TACL-ON and Cherry cells. Signals within the 3Mb region next to the TetO sites are excluded from this analysis. Peaks are divided into three groups, shared peaks between the two conditions, TACL-ON only, and Cherry-only. Note the similar levels between the two conditions across different genomic regions. **(C)** Average Hi-C loops (left), TAD boundaries (middle) and TADs (right) in TACL-ON, TACL-OFF, Cherry, and publicly available HAP1 cells (PMID: <u>26499245</u>, <u>28475897</u>). Value in the upper-right corners indicates the average loop strength, average TAD boundary strength, and average TAD density over the background, respectively. **(D)** Hi-C interaction matrices at 100-kb resolution showing examples of chromatin structure in TACL-ON, TACL-OFF, Cherry, and publicly available HAP1 cells (PMID: <u>26499245</u>, <u>28475897</u>) for Chr2: 135.0-165.0 Mb (left) and Chr21: 23.0–43.0 Mb (right). **(E)** Volcano plot depicting differentially expressed genes measured by BrU-seq in TACL-ON and Cherry cells. Pink dots represent genes expressed at higher levels in TACL-ON cells while green dots represent genes with higher expression levels in Cherry cells. Y-axis represents − log10 P values while X-axis shows log2 fold change values. **(F)** Measurement of cellular proliferation of TACL-ON and Cherry cells. Y-axis represents log2 cell number of each cell line measured.

**Figure S2.**
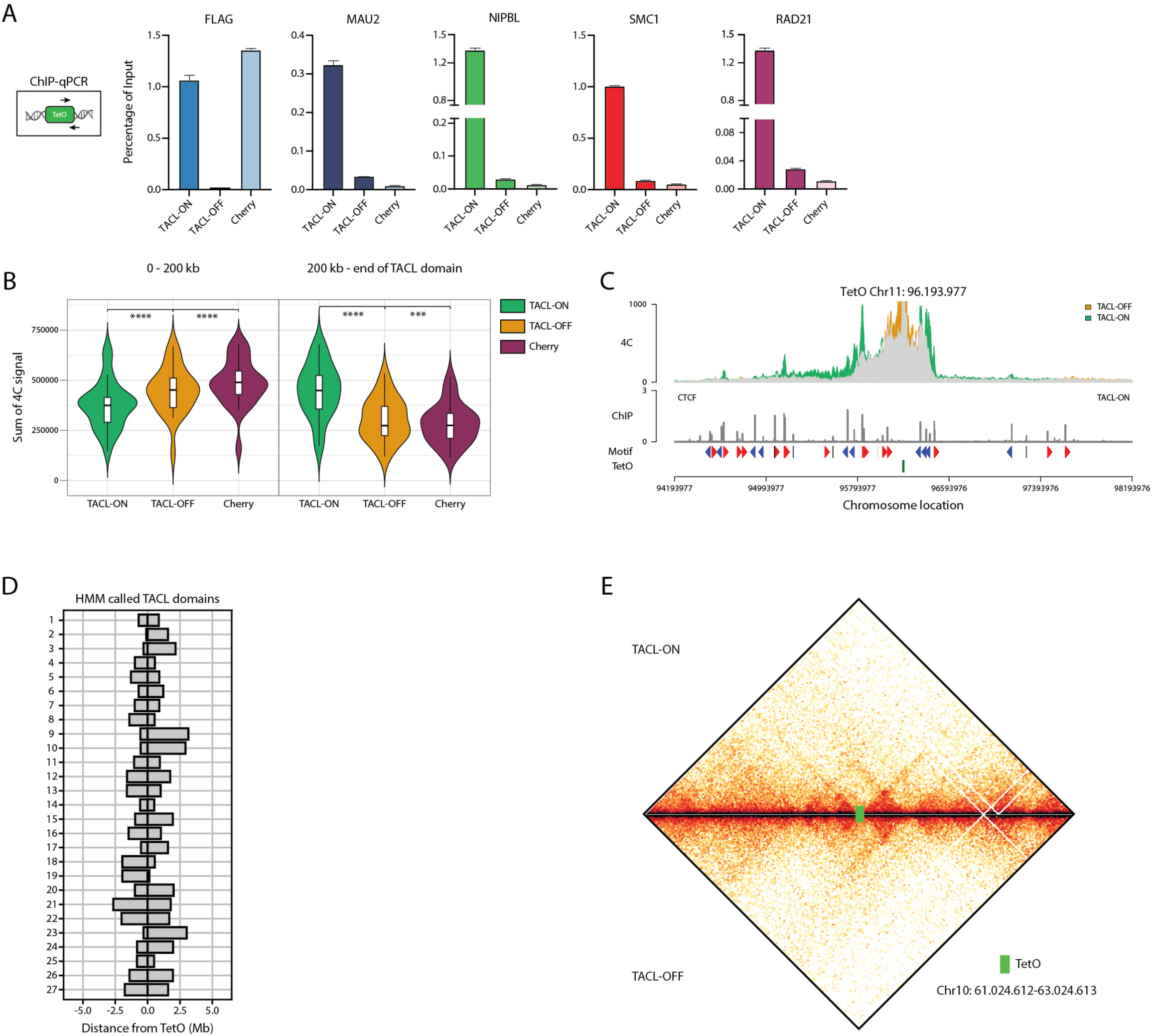
TACL efficiently recruits cohesin factors and induces topological changes on chromatin. **(A)** ChIP-qPCR analysis of FLAG, MAU2, NIPBL, SMC1, and RAD21 in TACL-ON, TACL-OFF, and Cherry cells. One set of primers was used to simultaneously detect all TetO platforms at once. Percentage of input was shown as y-axis. **(B)** Violin plots summarizing the 4C signals in TACL-ON, TACL-OFF, and Cherry cells. TetO platforms were used as viewpoint for the 4C experiments. 4C signals are divided into two groups, 0-200kb surrounding TetO platforms and 200-kb to the end of TACL domains. **** p value (paired t-test) < 0.0001; *** p value (paried t-test) <0.001. **(C)** An example 4C overlay of a TetO viewpoint on chr11. **(D)** HMM called TACL domains centred at the TetO platform integration locations. **(E)** An example locus of Hi-C analysis in TACL-ON and TACL-OFF cells. The location of TetO is highlighted in green. Note the stripes emerging from the TetO platform.

**Figure S3.**
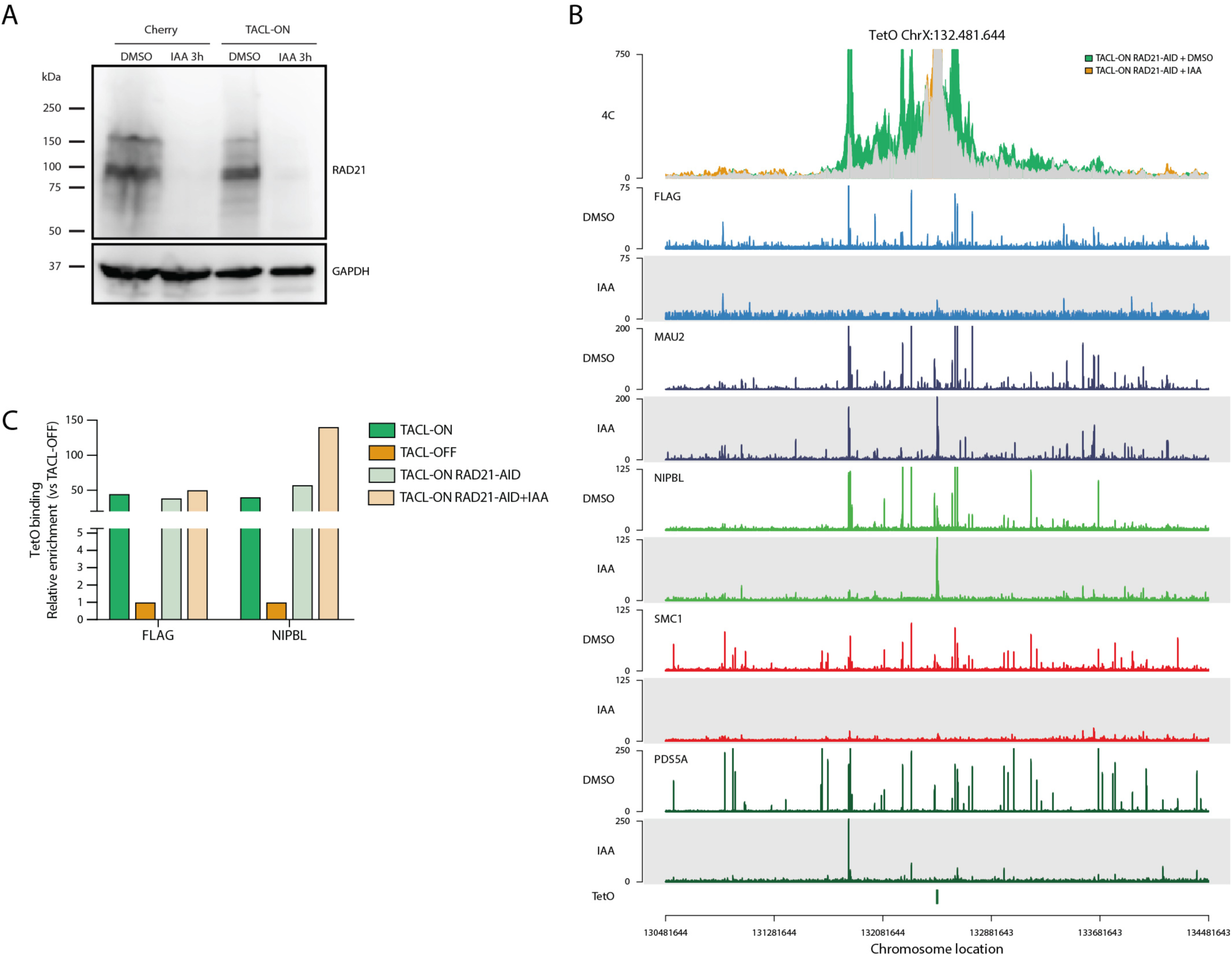
TACL-induced loop extrusion is RAD21 dependent. **(A)** Western blot analysis of RAD21 depletion in Cherry and TACL-ON cells. DMSO serves as a control and IAA treatment leads to RAD21 degradation. **(B)** An example TetO locus on chrX prensenting the 4C overlay and ChIP-seq tracks of FLAG, MAU2, NIPBL, SMC1, and PDS5A in RAD21-depleted cells. TACL-ON RAD21-AID+DMSO serves control. The plot is centered at the viewpoint of the 4C analysis (indicated as the TetO at the lower part of the panel). **(C)** Relative enrichment of FLAG and NIPBL at the TetO platforms. Values are normalized to TACL-OFF condition. Note the unchanged/increased binding of the factors in RAD21-depleted cells.

**Figure S4.**
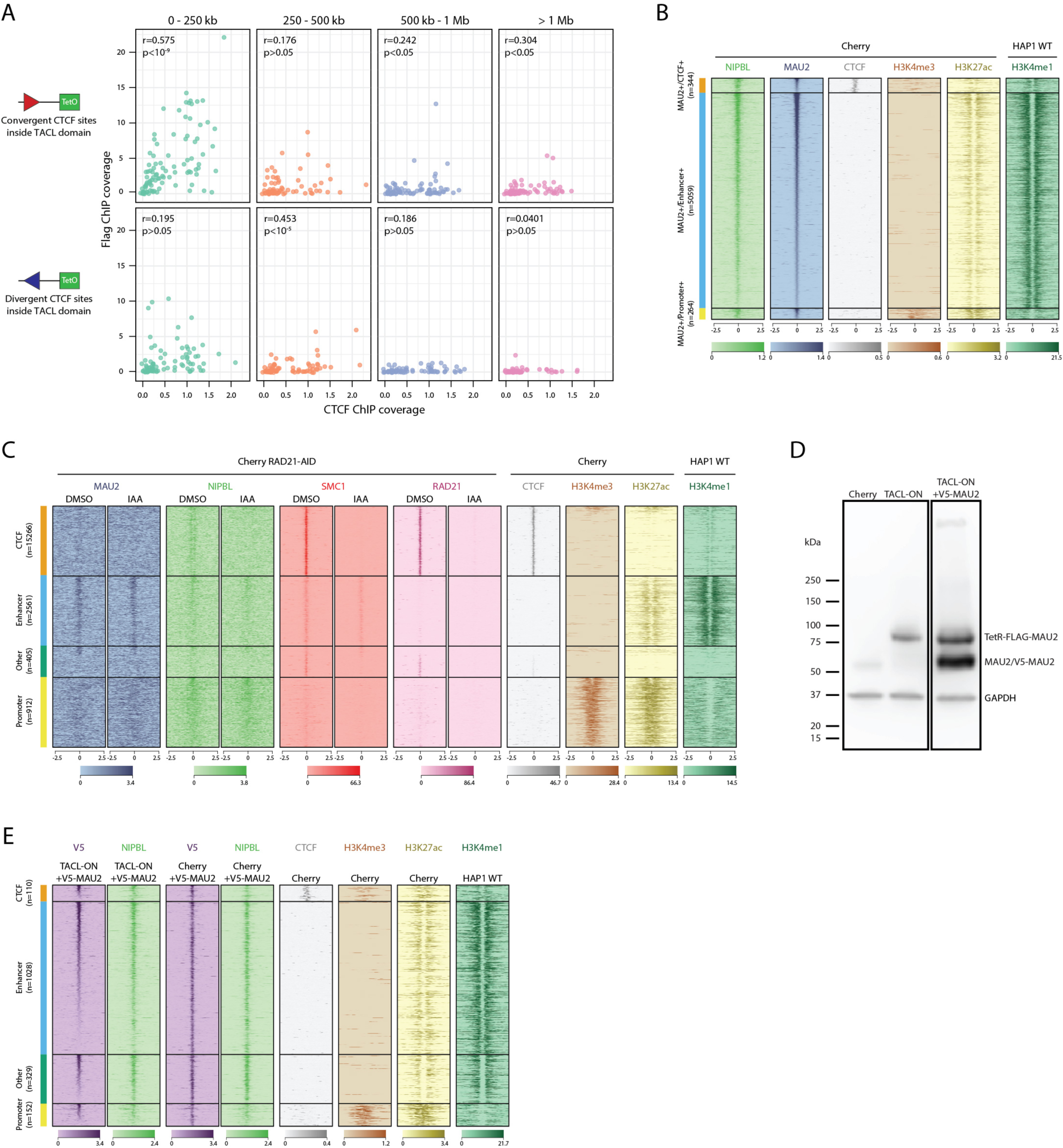
MAU2 is stably associated with cohesin from start to the end. **(A)** Correlation of FLAG and CTCF ChIP-seq signals. ChIP-seq signals are separated based on their distances to the TetO sites (cohesin loading sites). Convergent CTCF sites represents sites facing towards the TetO sites, while divergent CTCF sites represents sites facing away from the TetO sites. Pearson correlation coefficients (r) **and the** associated p-values are indicated” **(B)** Heatmaps of ChIP-seq signals of NIPBL, MAU2, CTCF, H3K4me3, and H3K27Ac in Cherry signals. H3K4me1 in WT HAP1 cells are taken as a reference. Genome-wide MAU2 ChIP-seq peaks in Cherry cells are divided into three groups based their colocalization with CTCF, enhancer (H3K4me3- and H3K4me1 or H3K27ac+), or promoter (H3K4me3+) histone marks. **(C)** Heatmaps of ChIP-seq signals for MAU2, NIPBL, SMC1, and RAD21 in Cherry RAD21-AID cells. For reference, CTCF, H3K4me3, H3K27Ac, and H3K4me1 signals are displayed. SMC1 and NIPBL peaks are categorized into four groups as indicated on the left, CTCF, Enhancer (H3K4me3- and H3K27Ac+ or H3K4me1+), Other (no active histone modification marks), and Promoter (H3K4me3+. ‘n’ stands for the number of peaks. **(D)** Western blot analysis of MAU2 expression in Cherry, TACL-ON, and TACL-ON+V5-MAU2 cells. Different versions of MAU2 are visualized on the same blot with MAU2 antibody. GAPDH serves as loading control. **(E)** Heatmaps of ChIP-seq signals for V5 (V5-MAU2) and NIPBL in TACL-ON and Cherry cells +/- V5-MAU2 co-expression. For reference, CTCF, H3K4me3, H3K27Ac, and H3K4me1 signals are displayed. V5 Peaks are classified into four groups as mentioned above.

**Figure S5.**
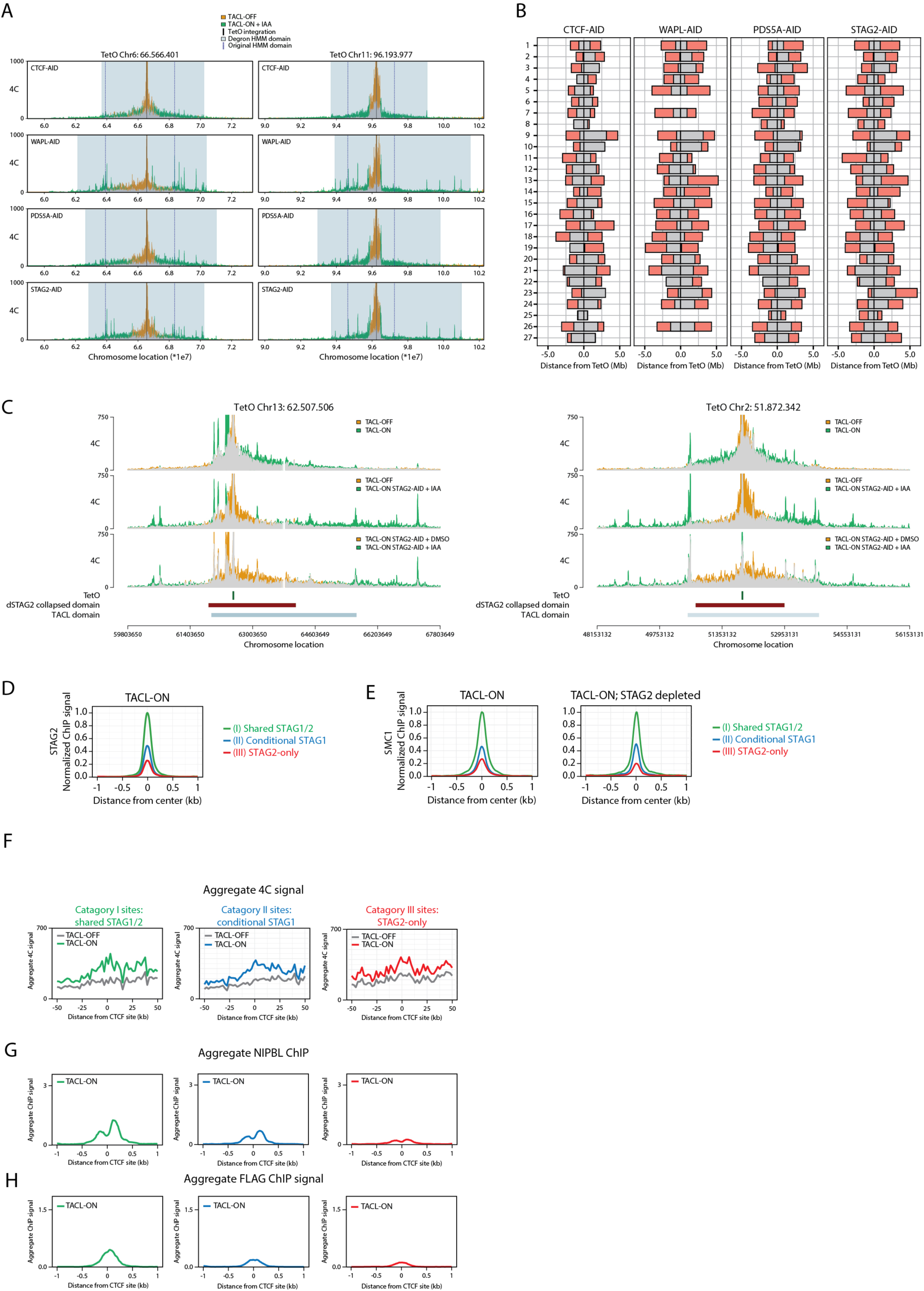
Depletion of cohesin factors cause extended loop extrusion. **(A)** 4C overlay at two example loci in different degron lines. Comparisons are made between TACL-OFF and TACL-ON, STAG2 depleted condition. 4C plots are centered around TetO sites. Original TACL domain are depicted by black lines and the extended domain after degron activation is depicted by grey squares. **(B)** Analysis of extended TACL-induced loop extrusion domain in different degron lines. Grey bars represent the original TACL domain in TACL-ON cells, while pink bars represent the extended TACL domain in depletion cells. **(C)** 4C overlay at two example loci. Comparisons are made between TACL-OFF and TACL-ON, TACL-OFF and TACL-ON, STAG2-AID+IAA, TACL-ON, STAG2-AID +DMSO and TACL-ON, STAG2-AID +IAA conditions. TACL domain and the collapsed intradomain after STAG2 depletion are depicted below. **(D)** Aggregate STAG2 ChIP signal in TACL-ON cells. Different STAG categories are depicted in the indicated colors. Average ChIP signals are normalized to the average signal at the convergent shared peaks_located inside the collapsed domains **(E)** Aggregate SMC1 ChIP signal in TACL-ON and TACL-ON, STAG2-depleted cells. Average ChIP signals are normalized to the average signal at the convergent shared peaks_located inside the collapsed domains **(F)** Aggregate 4C signal plots of TACL-ON and TACL-OFF cells within the STAG2-depleted collapsed domain. Only CTCF sites with TetO-divergent orientations are shown. Panels are divided by different STAG categories. TACL-OFF is indicated in grey and TACL-ON is indicated with the colors of each STAG category. Cat. I in green, cat. II in blue and cat. III in red. Signals are centered around CTCF binding sites. **(G)** Aggregate NIPBL ChIP signal at TetO-divergent CTCF sites in TACL-ON, STAG2-depleted cells. **(H)** Aggregate FLAG ChIP signal at TetO-divergent CTCF sites in TACL-ON, STAG2-depleted cells.

**Figure S6.**
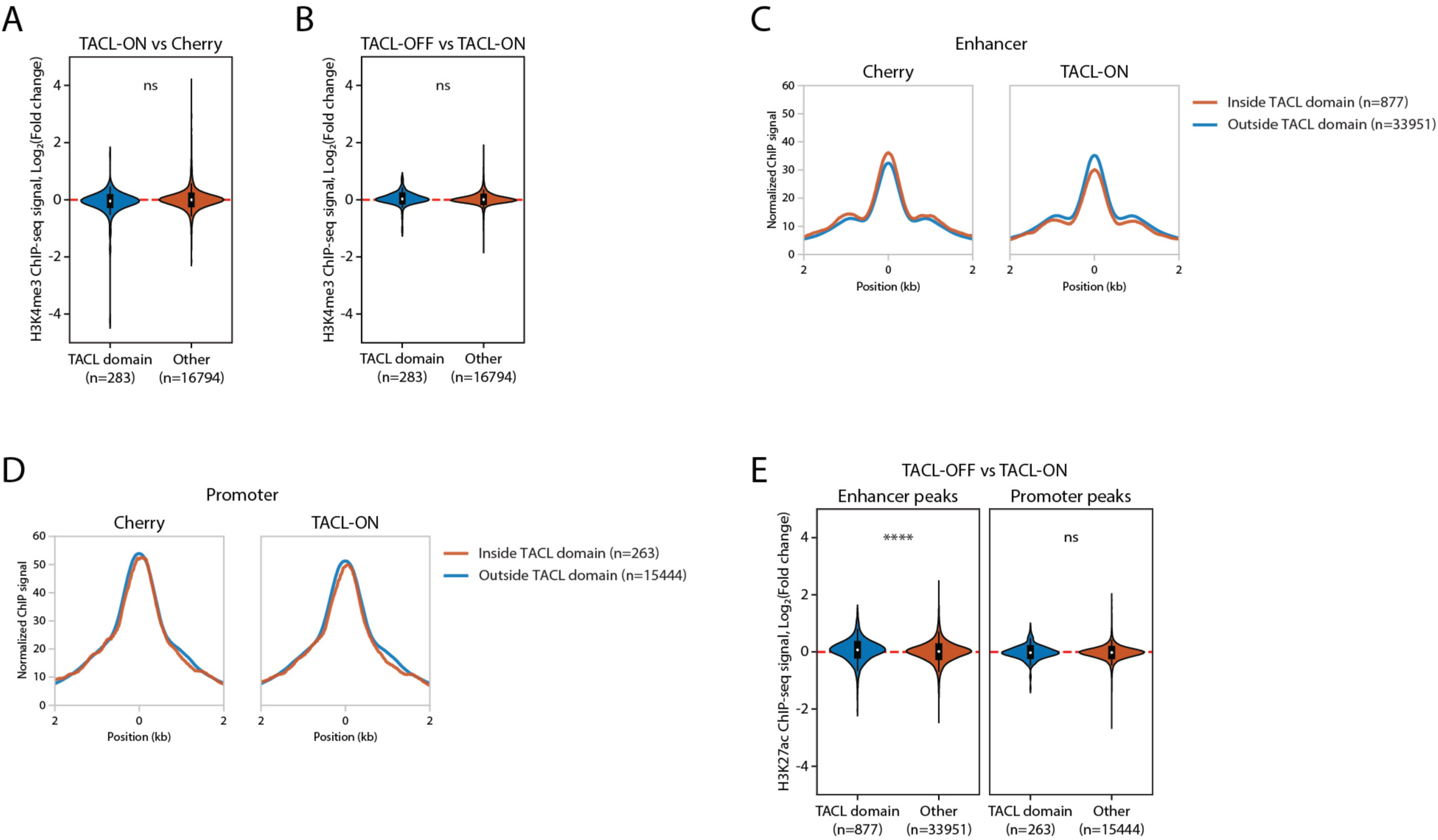
TACL-induced loop extrusion impact active histone modifications. **(A-B)** Average signal plot for H3K27Ac ChIP-seq signals at enhancers (A) and promoters (B). Comparisons are made between TACL-ON and Cherry cells. H3K27Ac peaks are divided into either inside TACL domain (red) or outside TACL domain. The number of peaks is depicted in the bracket of each category. ‘ns’ stands for not significant. **(C-D)** Violin plots showing H3K4me3 peaks in TACL-ON versus Cherry cells (C) or TACL-ON versus TACL-OFF cells (D). Y-axis are plotted as log2 fold change between the conditions. H3K4me3 peaks are divided into either inside or outside TACL domain. **(E)** Violin plots showing H3K27Ac peaks at enhancer and promoter sites in TACL-ON and TACL-OFF cells. Y-axis are plotted as log2 fold change between the conditions. H3K27Ac peaks are divided into either inside or outside TACL domain. **** p value < 0.0001; ‘ns’ stands for not significant.

## Notes

### Competing Interest Statement

The authors have declared no competing interest.

### Summary of Updates

The revised version includes new findings and insights in addition to our previous manuscript.

## References

1. Rao, S.S.P., Huang, S.C., Glenn St Hilaire, B., Engreitz, J.M., Perez, E.M., Kieffer-Kwon, K.R., Sanborn, A.L., Johnstone, S.E., Bascom, G.D., Bochkov, I.D., et al. (2017). Cohesin Loss Eliminates All Loop Domains. Cell 171, 305–320 e324. 10.1016/j.cell.2017.09.026.

2. Schwarzer, W., Abdennur, N., Goloborodko, A., Pekowska, A., Fudenberg, G., Loe-Mie, Y., Fonseca, N.A., Huber, W., Haering, C.H., Mirny, L., and Spitz, F. (2017). Two independent modes of chromatin organization revealed by cohesin removal. Nature 551, 51–56. 10.1038/nature24281.

3. Wutz, G., Varnai, C., Nagasaka, K., Cisneros, D.A., Stocsits, R.R., Tang, W., Schoenfelder, S., Jessberger, G., Muhar, M., Hossain, M.J., et al. (2017). Topologically associating domains and chromatin loops depend on cohesin and are regulated by CTCF, WAPL, and PDS5 proteins. EMBO J 36, 3573–3599. 10.15252/embj.201798004.

4. Fudenberg, G., Imakaev, M., Lu, C., Goloborodko, A., Abdennur, N., and Mirny, L.A. (2016). Formation of Chromosomal Domains by Loop Extrusion. Cell Rep 15, 2038–2049. 10.1016/j.celrep.2016.04.085.

5. Ganji, M., Shaltiel, I.A., Bisht, S., Kim, E., Kalichava, A., Haering, C.H., and Dekker, C. (2018). Real-time imaging of DNA loop extrusion by condensin. Science 360, 102–105. 10.1126/science.aar7831.

6. Sanborn, A.L., Rao, S.S., Huang, S.C., Durand, N.C., Huntley, M.H., Jewett, A.I., Bochkov, I.D., Chinnappan, D., Cutkosky, A., Li, J., et al. (2015). Chromatin extrusion explains key features of loop and domain formation in wild-type and engineered genomes. Proc Natl Acad Sci U S A 112, E6456–6465. 10.1073/pnas.1518552112.

7. Datta, S., Lecomte, L., and Haering, C.H. (2020). Structural insights into DNA loop extrusion by SMC protein complexes. Curr Opin Struct Biol 65, 102–109. 10.1016/j.sbi.2020.06.009.

8. Davidson, I.F., and Peters, J.M. (2021). Genome folding through loop extrusion by SMC complexes. Nat Rev Mol Cell Biol 22, 445–464. 10.1038/s41580-021-00349-7.

9. Hoencamp, C., and Rowland, B.D. (2023). Genome control by SMC complexes. Nat Rev Mol Cell Biol 24, 633–650. 10.1038/s41580-023-00609-8.

10. Mirny, L., and Dekker, J. (2022). Mechanisms of Chromosome Folding and Nuclear Organization: Their Interplay and Open Questions. Cold Spring Harb Perspect Biol 14. 10.1101/cshperspect.a040147.

11. Kojic, A., Cuadrado, A., De Koninck, M., Gimenez-Llorente, D., Rodriguez-Corsino, M., Gomez-Lopez, G., Le Dily, F., Marti-Renom, M.A., and Losada, A. (2018). Distinct roles of cohesin-SA1 and cohesin-SA2 in 3D chromosome organization. Nat Struct Mol Biol 25, 496–504. 10.1038/s41594-018-0070-4.

12. Cuadrado, A., Gimenez-Llorente, D., Kojic, A., Rodriguez-Corsino, M., Cuartero, Y., Martin-Serrano, G., Gomez-Lopez, G., Marti-Renom, M.A., and Losada, A. (2019). Specific Contributions of Cohesin-SA1 and Cohesin-SA2 to TADs and Polycomb Domains in Embryonic Stem Cells. Cell Rep 27, 3500–3510 e3504. 10.1016/j.celrep.2019.05.078.

13. van der Weide, R.H., van den Brand, T., Haarhuis, J.H.I., Teunissen, H., Rowland, B.D., and de Wit, E. (2021). Hi-C analyses with GENOVA: a case study with cohesin variants. NAR Genom Bioinform 3, lqab040. 10.1093/nargab/lqab040.

14. Liu, N.Q., Maresca, M., van den Brand, T., Braccioli, L., Schijns, M., Teunissen, H., Bruneau, B.G., Nora, E.P., and de Wit, E. (2021). WAPL maintains a cohesin loading cycle to preserve cell-type-specific distal gene regulation. Nat Genet 53, 100–109. 10.1038/s41588-020-00744-4.

15. Vian, L., Pekowska, A., Rao, S.S.P., Kieffer-Kwon, K.R., Jung, S., Baranello, L., Huang, S.C., El Khattabi, L., Dose, M., Pruett, N., et al. (2018). The Energetics and Physiological Impact of Cohesin Extrusion. Cell 175, 292–294. 10.1016/j.cell.2018.09.002.

16. Ciosk, R., Shirayama, M., Shevchenko, A., Tanaka, T., Toth, A., Shevchenko, A., and Nasmyth, K. (2000). Cohesin’s binding to chromosomes depends on a separate complex consisting of Scc2 and Scc4 proteins. Mol Cell 5, 243–254. 10.1016/s1097-2765(00)80420-7.

17. Petela, N.J., Gonzalez Llamazares, A., Dixon, S., Hu, B., Lee, B.G., Metson, J., Seo, H., Ferrer-Harding, A., Voulgaris, M., Gligoris, T., et al. (2021). Folding of cohesin’s coiled coil is important for Scc2/4-induced association with chromosomes. Elife 10. 10.7554/eLife.67268.

18. Bauer, B.W., Davidson, I.F., Canena, D., Wutz, G., Tang, W., Litos, G., Horn, S., Hinterdorfer, P., and Peters, J.M. (2021). Cohesin mediates DNA loop extrusion by a “swing and clamp” mechanism. Cell 184, 5448–5464 e5422. 10.1016/j.cell.2021.09.016.

19. Davidson, I.F., Bauer, B., Goetz, D., Tang, W., Wutz, G., and Peters, J.M. (2019). DNA loop extrusion by human cohesin. Science 366, 1338–1345. 10.1126/science.aaz3418.

20. Kim, Y., Shi, Z., Zhang, H., Finkelstein, I.J., and Yu, H. (2019). Human cohesin compacts DNA by loop extrusion. Science 366, 1345–1349. 10.1126/science.aaz4475.

21. Bastie, N., Chapard, C., Dauban, L., Gadal, O., Beckouet, F., and Koszul, R. (2022). Smc3 acetylation, Pds5 and Scc2 control the translocase activity that establishes cohesin-dependent chromatin loops. Nat Struct Mol Biol 29, 575–585. 10.1038/s41594-022-00780-0.

22. Haarhuis, J.H.I., van der Weide, R.H., Blomen, V.A., Yanez-Cuna, J.O., Amendola, M., van Ruiten, M.S., Krijger, P.H.L., Teunissen, H., Medema, R.H., van Steensel, B., et al. (2017). The Cohesin Release Factor WAPL Restricts Chromatin Loop Extension. Cell 169, 693–707.e614. 10.1016/j.cell.2017.04.013.

23. de Wit, E., Vos, E.S., Holwerda, S.J., Valdes-Quezada, C., Verstegen, M.J., Teunissen, H., Splinter, E., Wijchers, P.J., Krijger, P.H., and de Laat, W. (2015). CTCF Binding Polarity Determines Chromatin Looping. Mol Cell 60, 676–684. 10.1016/j.molcel.2015.09.023.

24. Li, Y., Haarhuis, J.H.I., Sedeno Cacciatore, A., Oldenkamp, R., van Ruiten, M.S., Willems, L., Teunissen, H., Muir, K.W., de Wit, E., Rowland, B.D., and Panne, D. (2020). The structural basis for cohesin-CTCF-anchored loops. Nature 578, 472–476. 10.1038/s41586-019-1910-z.

25. Rao, S.S., Huntley, M.H., Durand, N.C., Stamenova, E.K., Bochkov, I.D., Robinson, J.T., Sanborn, A.L., Machol, I., Omer, A.D., Lander, E.S., and Aiden, E.L. (2014). A 3D map of the human genome at kilobase resolution reveals principles of chromatin looping. Cell 159, 1665–1680. 10.1016/j.cell.2014.11.021.

26. Yu, D., Chen, G., Wang, Y., Wang, Y., Lin, R., Liu, N., Zhu, P., Liu, H., Hu, T., Feng, R., et al. (2022). Regulation of cohesin-mediated chromosome folding by PDS5 in mammals. EMBO Rep 23, e54853. 10.15252/embr.202254853.

27. Nora, E.P., Caccianini, L., Fudenberg, G., So, K., Kameswaran, V., Nagle, A., Uebersohn, A., Hajj, B., Saux, A.L., Coulon, A., et al. (2020). Molecular basis of CTCF binding polarity in genome folding. Nat Commun 11, 5612. 10.1038/s41467-020-19283-x.

28. Petela, N.J., Gligoris, T.G., Metson, J., Lee, B.G., Voulgaris, M., Hu, B., Kikuchi, S., Chapard, C., Chen, W., Rajendra, E., et al. (2018). Scc2 Is a Potent Activator of Cohesin’s ATPase that Promotes Loading by Binding Scc1 without Pds5. Mol Cell 70, 1134–1148 e1137. 10.1016/j.molcel.2018.05.022.

29. Cuadrado, A., Gimenez-Llorente, D., De Koninck, M., Ruiz-Torres, M., Kojic, A., Rodriguez-Corsino, M., and Losada, A. (2022). Contribution of variant subunits and associated factors to genome-wide distribution and dynamics of cohesin. Epigenetics Chromatin 15, 37. 10.1186/s13072-022-00469-0.

30. Dauban, L., Montagne, R., Thierry, A., Lazar-Stefanita, L., Bastie, N., Gadal, O., Cournac, A., Koszul, R., and Beckouet, F. (2020). Regulation of Cohesin-Mediated Chromosome Folding by Eco1 and Other Partners. Mol Cell 77, 1279–1293 e1274. 10.1016/j.molcel.2020.01.019.

31. Banigan, E.J., Tang, W., van den Berg, A.A., Stocsits, R.R., Wutz, G., Brandao, H.B., Busslinger, G.A., Peters, J.M., and Mirny, L.A. (2023). Transcription shapes 3D chromatin organization by interacting with loop extrusion. Proc Natl Acad Sci U S A 120, e2210480120. 10.1073/pnas.2210480120.

32. Sexton, T., Platania, A., Erb, C., Barbieri, M., Molcrette, B., Grandgirard, E., de Kort, M., Meabum, K., Taylor, T., Shchuka, V., et al. (2023). Competition between transcription and loop extrusion modulates promoter and enhancer dynamics. Res Sq. 10.21203/rs.3.rs-3164817/v1.

33. Zhang, S., Ubelmesser, N., Barbieri, M., and Papantonis, A. (2023). Enhancer-promoter contact formation requires RNAPII and antagonizes loop extrusion. Nat Genet 55, 832–840. 10.1038/s41588-023-01364-4.

34. Calderon, L., Weiss, F.D., Beagan, J.A., Oliveira, M.S., Georgieva, R., Wang, Y.F., Carroll, T.S., Dharmalingam, G., Gong, W., Tossell, K., et al. (2022). Cohesin-dependence of neuronal gene expression relates to chromatin loop length. Elife 11. 10.7554/eLife.76539.

35. Kane, L., Williamson, I., Flyamer, I.M., Kumar, Y., Hill, R.E., Lettice, L.A., and Bickmore, W.A. (2022). Cohesin is required for long-range enhancer action at the Shh locus. Nat Struct Mol Biol 29, 891–897. 10.1038/s41594-022-00821-8.

36. Rinzema, N.J., Sofiadis, K., Tjalsma, S.J.D., Verstegen, M., Oz, Y., Valdes-Quezada, C., Felder, A.K., Filipovska, T., van der Elst, S., de Andrade Dos Ramos, Z., et al. (2022). Building regulatory landscapes reveals that an enhancer can recruit cohesin to create contact domains, engage CTCF sites and activate distant genes. Nat Struct Mol Biol 29, 563–574. 10.1038/s41594-022-00787-7.

37. Thiecke, M.J., Wutz, G., Muhar, M., Tang, W., Bevan, S., Malysheva, V., Stocsits, R., Neumann, T., Zuber, J., Fraser, P., et al. (2020). Cohesin-Dependent and -Independent Mechanisms Mediate Chromosomal Contacts between Promoters and Enhancers. Cell Rep 32, 107929. 10.1016/j.celrep.2020.107929.

38. Gabriele, M., Brandao, H.B., Grosse-Holz, S., Jha, A., Dailey, G.M., Cattoglio, C., Hsieh, T.S., Mirny, L., Zechner, C., and Hansen, A.S. (2022). Dynamics of CTCF- and cohesin-mediated chromatin looping revealed by live-cell imaging. Science 376, 496–501. 10.1126/science.abn6583.

39. Redolfi, J., Zhan, Y., Valdes-Quezada, C., Kryzhanovska, M., Guerreiro, I., Iesmantavicius, V., Pollex, T., Grand, R.S., Mulugeta, E., Kind, J., et al. (2019). DamC reveals principles of chromatin folding in vivo without crosslinking and ligation. Nat Struct Mol Biol 26, 471–480. 10.1038/s41594-019-0231-0.

40. Hinshaw, S.M., Makrantoni, V., Kerr, A., Marston, A.L., and Harrison, S.C. (2015). Structural evidence for Scc4-dependent localization of cohesin loading. Elife 4, e06057. 10.7554/eLife.06057.

41. Watrin, E., Schleiffer, A., Tanaka, K., Eisenhaber, F., Nasmyth, K., and Peters, J.M. (2006). Human Scc4 is required for cohesin binding to chromatin, sister-chromatid cohesion, and mitotic progression. Curr Biol 16, 863–874. 10.1016/j.cub.2006.03.049.

42. Guo, Y., Al-Jibury, E., Garcia-Millan, R., Ntagiantas, K., King, J.W.D., Nash, A.J., Galjart, N., Lenhard, B., Rueckert, D., Fisher, A.G., et al. (2022). Chromatin jets define the properties of cohesin-driven in vivo loop extrusion. Mol Cell 82, 3769–3780 e3765. 10.1016/j.molcel.2022.09.003.

43. Vietri Rudan, M., Barrington, C., Henderson, S., Ernst, C., Odom, D.T., Tanay, A., and Hadjur, S. (2015). Comparative Hi-C reveals that CTCF underlies evolution of chromosomal domain architecture. Cell Rep 10, 1297–1309. 10.1016/j.celrep.2015.02.004.

44. Yesbolatova, A., Saito, Y., Kitamoto, N., Makino-Itou, H., Ajima, R., Nakano, R., Nakaoka, H., Fukui, K., Gamo, K., Tominari, Y., et al. (2020). The auxin-inducible degron 2 technology provides sharp degradation control in yeast, mammalian cells, and mice. Nat Commun 11, 5701. 10.1038/s41467-020-19532-z.

45. Kim, E., Kerssemakers, J., Shaltiel, I.A., Haering, C.H., and Dekker, C. (2020). DNA-loop extruding condensin complexes can traverse one another. Nature 579, 438–442. 10.1038/s41586-020-2067-5.

46. Allahyar, A., Vermeulen, C., Bouwman, B.A.M., Krijger, P.H.L., Verstegen, M., Geeven, G., van Kranenburg, M., Pieterse, M., Straver, R., Haarhuis, J.H.I., et al. (2018). Enhancer hubs and loop collisions identified from single-allele topologies. Nat Genet 50, 1151–1160. 10.1038/s41588-018-0161-5.

47. Chang, L.H., Ghosh, S., Papale, A., Luppino, J.M., Miranda, M., Piras, V., Degrouard, J., Edouard, J., Poncelet, M., Lecouvreur, N., et al. (2023). Multi-feature clustering of CTCF binding creates robustness for loop extrusion blocking and Topologically Associating Domain boundaries. Nat Commun 14, 5615. 10.1038/s41467-023-41265-y.

48. Hung, T.C., Kingsley, D.M., and Boettiger, A.N. (2024). Boundary stacking interactions enable cross-TAD enhancer-promoter communication during limb development. Nat Genet 56, 306–314. 10.1038/s41588-023-01641-2.

49. van Ruiten, M.S., van Gent, D., Sedeno Cacciatore, A., Fauster, A., Willems, L., Hekkelman, M.L., Hoekman, L., Altelaar, M., Haarhuis, J.H.I., Brummelkamp, T.R., et al. (2022). The cohesin acetylation cycle controls chromatin loop length through a PDS5A brake mechanism. Nat Struct Mol Biol 29, 586–591. 10.1038/s41594-022-00773-z.

50. Nora, E.P., Goloborodko, A., Valton, A.L., Gibcus, J.H., Uebersohn, A., Abdennur, N., Dekker, J., Mirny, L.A., and Bruneau, B.G. (2017). Targeted Degradation of CTCF Decouples Local Insulation of Chromosome Domains from Genomic Compartmentalization. Cell 169, 930–944 e922. 10.1016/j.cell.2017.05.004.

51. Galvan, D.L., Nakazawa, Y., Kaja, A., Kettlun, C., Cooper, L.J., Rooney, C.M., and Wilson, M.H. (2009). Genome-wide mapping of PiggyBac transposon integrations in primary human T cells. J Immunother 32, 837–844. 10.1097/CJI.0b013e3181b2914c.

52. Holzmann, J., Politi, A.Z., Nagasaka, K., Hantsche-Grininger, M., Walther, N., Koch, B., Fuchs, J., Durnberger, G., Tang, W., Ladurner, R., et al. (2019). Absolute quantification of cohesin, CTCF and their regulators in human cells. Elife 8. 10.7554/eLife.46269.

53. Wutz, G., Ladurner, R., St Hilaire, B.G., Stocsits, R.R., Nagasaka, K., Pignard, B., Sanborn, A., Tang, W., Varnai, C., Ivanov, M.P., et al. (2020). ESCO1 and CTCF enable formation of long chromatin loops by protecting cohesin(STAG1) from WAPL. Elife 9. 10.7554/eLife.52091.

54. Krijger, P.H.L., Geeven, G., Bianchi, V., Hilvering, C.R.E., and de Laat, W. (2020). 4C-seq from beginning to end: A detailed protocol for sample preparation and data analysis. Methods 170, 17–32. 10.1016/j.ymeth.2019.07.014.

55. van de Werken, H.J., Landan, G., Holwerda, S.J., Hoichman, M., Klous, P., Chachik, R., Splinter, E., Valdes-Quezada, C., Oz, Y., Bouwman, B.A., et al. (2012). Robust 4C-seq data analysis to screen for regulatory DNA interactions. Nat Methods 9, 969–972. 10.1038/nmeth.2173.

56. Vermeulen, C., Allahyar, A., Bouwman, B.A.M., Krijger, P.H.L., Verstegen, M., Geeven, G., Valdes-Quezada, C., Renkens, I., Straver, R., Kloosterman, W.P., et al. (2020). Multi-contact 4C: long-molecule sequencing of complex proximity ligation products to uncover local cooperative and competitive chromatin topologies. Nat Protoc 15, 364–397. 10.1038/s41596-019-0242-7.

57. Roberts, T.C., Hart, J.R., Kaikkonen, M.U., Weinberg, M.S., Vogt, P.K., and Morris, K.V. (2015). Quantification of nascent transcription by bromouridine immunocapture nuclear run-on RT-qPCR. Nat Protoc 10, 1198–1211. 10.1038/nprot.2015.076.

58. Lawrence, M., Gentleman, R., and Carey, V. (2009). rtracklayer: an R package for interfacing with genome browsers. Bioinformatics 25, 1841–1842. 10.1093/bioinformatics/btp328.

59. Lawrence, M., Huber, W., Pages, H., Aboyoun, P., Carlson, M., Gentleman, R., Morgan, M.T., and Carey, V.J. (2013). Software for computing and annotating genomic ranges. PLoS Comput Biol 9, e1003118. 10.1371/journal.pcbi.1003118.

60. Grant, C.E., Bailey, T.L., and Noble, W.S. (2011). FIMO: scanning for occurrences of a given motif. Bioinformatics 27, 1017–1018. 10.1093/bioinformatics/btr064.

61. Khan, A., Fornes, O., Stigliani, A., Gheorghe, M., Castro-Mondragon, J.A., van der Lee, R., Bessy, A., Cheneby, J., Kulkarni, S.R., Tan, G., et al. (2018). JASPAR 2018: update of the open-access database of transcription factor binding profiles and its web framework. Nucleic Acids Res 46, D1284. 10.1093/nar/gkx1188.

